# Molecular Tuning of Styryl Dyes Leads to Versatile and Efficient Plasma Membrane Probes for Cell and Tissue Imaging

**DOI:** 10.1101/819383

**Authors:** Mayeul Collot, Emmanuel Boutant, Kyong Tkhe Fam, Lydia Danglot, Andrey S. Klymchenko

## Abstract

The plasma membrane (PM) plays a major role in many biological processes; therefore its proper fluorescence staining is required in bioimaging. Among the commercially available PM probes, styryl dye FM1-43 is one of the most widely used. In this work, we demonstrated that fine chemical modifications of FM1-43 can dramatically improve the PM staining. The newly developed probes, SP-468 and SQ-535 were found to display enhanced photophysical properties (reduced crosstalk, higher brightness, improved photostability) and unlike FM1-43, provided excellent and immediate PM staining in 5 different mammalian cell lines including neurons (primary culture and tissue imaging). Additionally, we showed that the new probes displayed differences in their internalization pathways compared to their parent FM1-43. Finally, we demonstrated that the modifications made to FM1-43 did not impair the ability of the new probes to stain the PM of plant cells. Overall, this work presents new useful probes for PM imaging in cells and tissues and provides insights on the molecular design of new PM targeting molecules.

## Introduction

The plasma membrane (PM), as a barrier between the extracellular and intracellular environments, maintains the integrity of various organisms at the cellular level and plays key roles in biology.^1, 2 3^ Consequently, the proper and selective staining of the PM is important in bioimaging to delimit the cells and to continue deciphering its role in various cellular mechanisms. Nowadays, fluorescence microscopy remains the most widely used bioimaging technique as it is relatively affordable and it rapidly provides important information with good resolution. From these two observations, numerous efforts have been made to develop fluorescent PM probes^4, 5, 6, 7^ including from our group.^8-15^ Among them, FM dyes (named after Fei Mao^16^), which are styryl dyes introduced by Betz and colleagues in the early 90s’ as synaptic vesicle markers,^16^ rapidly became among the most famous and used commercially available PM probes.^17, 18^ These amphiphilic dyes were further developed by the group of Loew to give rise to PM voltage sensitive probes^19, 20^ and membrane lipid domains probes.^21^ These probes are composed of an aniline or naphthylamine donor linked to a pyridinium acceptor moiety through a double bond or a polymethine chain, which can tune their color.^19, 20^ However, efforts were mainly focused on designing probes bearing polar groups on one end and hydrophobic groups on the other end, which thus imposed vertical orientation of styryl fluorophores in the membrane. In the case of FM1-43 (Figure 1), one of the most commonly used FM probes, it was claimed that the dibutyl chains bore by the aniline moiety interact with the hydrocarbon chains of the lipid bilayer via hydrophobic interactions whereas the triethylammonium cationic moiety bore by the pyridinium interacts with the negative charges of the phospholipids through opposite charge attraction.^22^ Unlike hydrophobic carbocyanines, such as DiI, DiD and DiR, bearing two C-18 hydrocarbon chains,^17^ FM dyes display a lower hydrophobicity and thus do not precipitate in aqueous media. Therefore, FM probes are successfully and widely used for imaging PM or vesicle trafficking in plants.^23, 24^ In mammalian cells they were applied to study vesiculation^25, 26^ and to image recycled synaptic vesicles in neuroscience.^27, 28, 29^ Unfortunately, the reduced hydrophobicity of FM1-43 leads to a lack of affinity towards the PM and inefficient PM staining in fixed cells.^4^ Moreover, FM probes tend to internalize quickly by endocytosis^30^ and their broad absorption and emission spectra limit their use in multicolor imaging. Nevertheless, attractive optical properties of FM dyes, such as their photostability, can constitute an advantage for long-term imaging and their large Stokes shifts is important Stimulated Emission Depletion (STED) super resolution imaging.^31, 32^ We recently introduced a family of PM probes with balanced hydrophobicity that provided efficient PM staining for cells and tissue, namely MemBright.^9, 14^ These PM probes were obtained by decorating BODIPY^9^ and cyanines^14^ with a clickable and selective PM targeting moiety: CAZ^10^ (Scheme 1, Figure 1).

**Figure 1.**
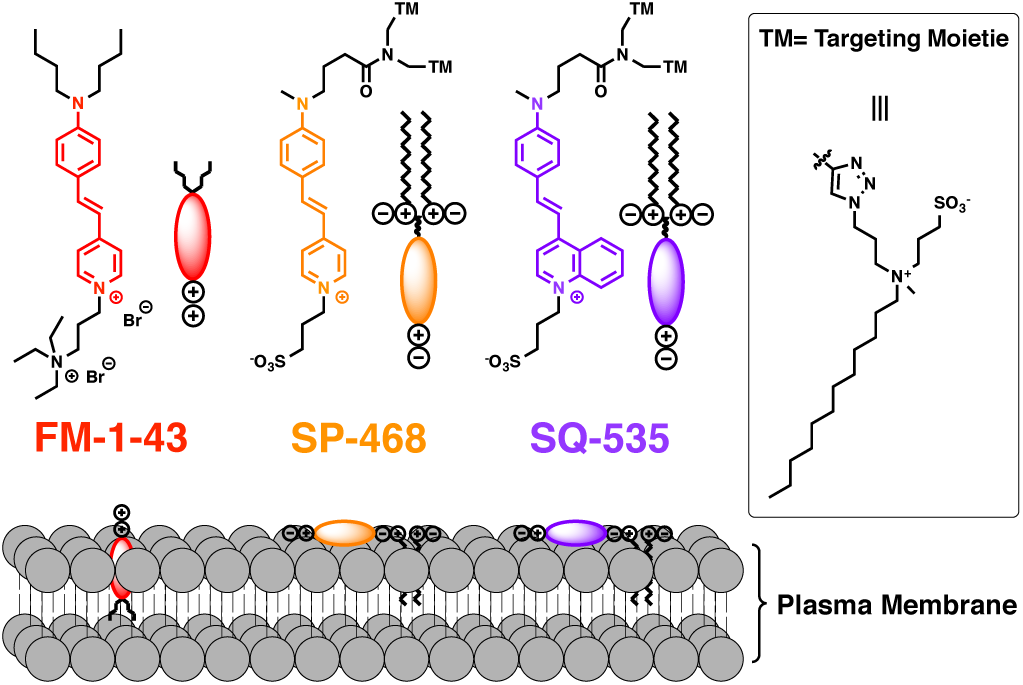
Structures of FM1-43 and the newly developed styryl probes SP-468 (Styryl Pyridinium - λ_Ex max_ 468) and SQ-535 (Styryl Quinolinium - λ_Ex max_ 535) and their orientation within the plasma membrane.

We herein showed that the staining properties of FM dyes can be dramatically improved by means of fine chemical modifications. By neutralizing the global charge, inserting polar head groups at both ends of the probe, tuning the hydrophobicity and optionally extending the electronic system of the dye, we obtained two new optimized version of FM1-43, namely SP-468 and SQ-535 (Figure 1) that efficiently and rapidly stain the PM in both eukaryotic and plant cells.

## Results and discussion

### Design of the probes

FM1-43 is a red emitting plasma membrane probe from the FM family. It is a styryl dye composed of an *N,N*-dialkylated aniline donor moiety linked through an ethylene connector to a 4-pyridinium acceptor moiety (Figure 1). It exhibits some affinity to PM due to two synergic interactions. First, the butyl groups bring hydrophobicity to enhance the affinity towards the hydrophobic tails of the lipids, while the cationic pyridinium and the ammonium moieties interact with the polar head of the lipids.^22^ In order to improve the PM imaging properties of FM1-43, we performed three different modifications. 1) We first chose to annihilate the cationic nature of FM dyes since cationic and hydrophobic molecules tend to accumulate inside the cells.^41^ Therefore, the ammonium group of FM was replaced by a sulfonate group to give rise to a neutral zwiterionic dye (Figure 1). 2) Then, one of the two butyl groups borne by the aniline moiety of FM1-43 was replaced by a methyl group in order to reduce the inductive donor effect that electronically enriches the nitrogen. This modification should provoke a hypsochromic shift in the absorption spectrum and thus should reduce the cross talk phenomenon due to the relatively high extinction coefficient of FM1-43 at 530 nm and 560 nm that are generally dedicated to excitation sources for the red channel in fluorescence microscopy. 3) Finally, two amphiphilic moieties were introduced by CuAAc click chemistry to ensure an efficient PM targeting.^9^ All these modifications led to a probe called SP-468 (Styryl Pyridinium). As an additional modification, the pyridinium moiety of the fluorophore has been replaced by a quinolinium in order to extend the π-conjugation of the fluorophore.^42^ This extension led to SQ-535 and is expected to provoke a bathochromic shift in both absorbance and emission spectra, which would ensure its use in the red channel with limited crosstalk in the green one.

### Synthesis

**Scheme 1.**
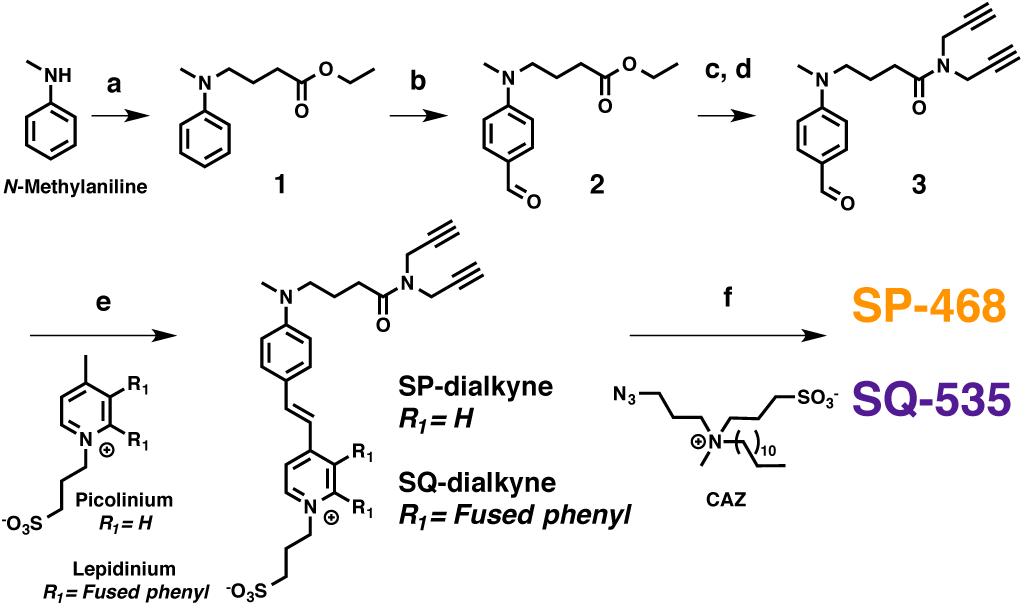
Synthesis of SP-468 and SQ-535. a) Ethyl 4-bromobutyrate, NaI, NaHCO_3_, DMSO, 80°C. b) POCl_3_, DMF, 55°C. c) NaOH (1M), MeOH. d) Dipropargylamine, DIEA, HATU, DMF, RT. e) Picolinium propane-1-sulfonate or lepidinium propane-1-sulfonate, piperidine (cat.), MeOH, 110°C. f) CAZ, CuSO_4_·5H_2_O, AscNa, water, DMF, 60°C.

The synthesis started with the *N*-alkylation of *N*-methylaniline to introduce an ester group (Scheme 1). The aniline 1 was then formylated using the Vilsmeier reaction to obtain 2. The ester was saponified and the obtained carboxylic acid was directly coupled to dipropargylamine using HATU as a coupling agent to give compound 3. This aldehyde was then involved in Knoevenagel reaction with picolinium propane-1-sulfonate and lepidinium propane-1-sulfonate to obtain SP-dialkyne and SQ-dialkyne respectively. These styryl dyes were clicked to CAZ^10^ to finally lead to the probes SP-468 and SQ-535 (Scheme 1).

### Spectroscopic studies

To evaluate their photophysical properties, the absorption and emission spectra of the three styryl dyes were measured in various conditions. Methanol and DMSO were chosen as polar and solubilizing solvents, whereas PBS was chosen as a typical aqueous medium for cell imaging microscopy. In order to mimic the presence of plasma membrane, liposomes composed of DOPC (1,2-Dioleoyl-*sn*-glycero-3-phosphocholine) were used in the presence of the dyes. Their photophysical properties are reported in table 1.

**Table 1.**
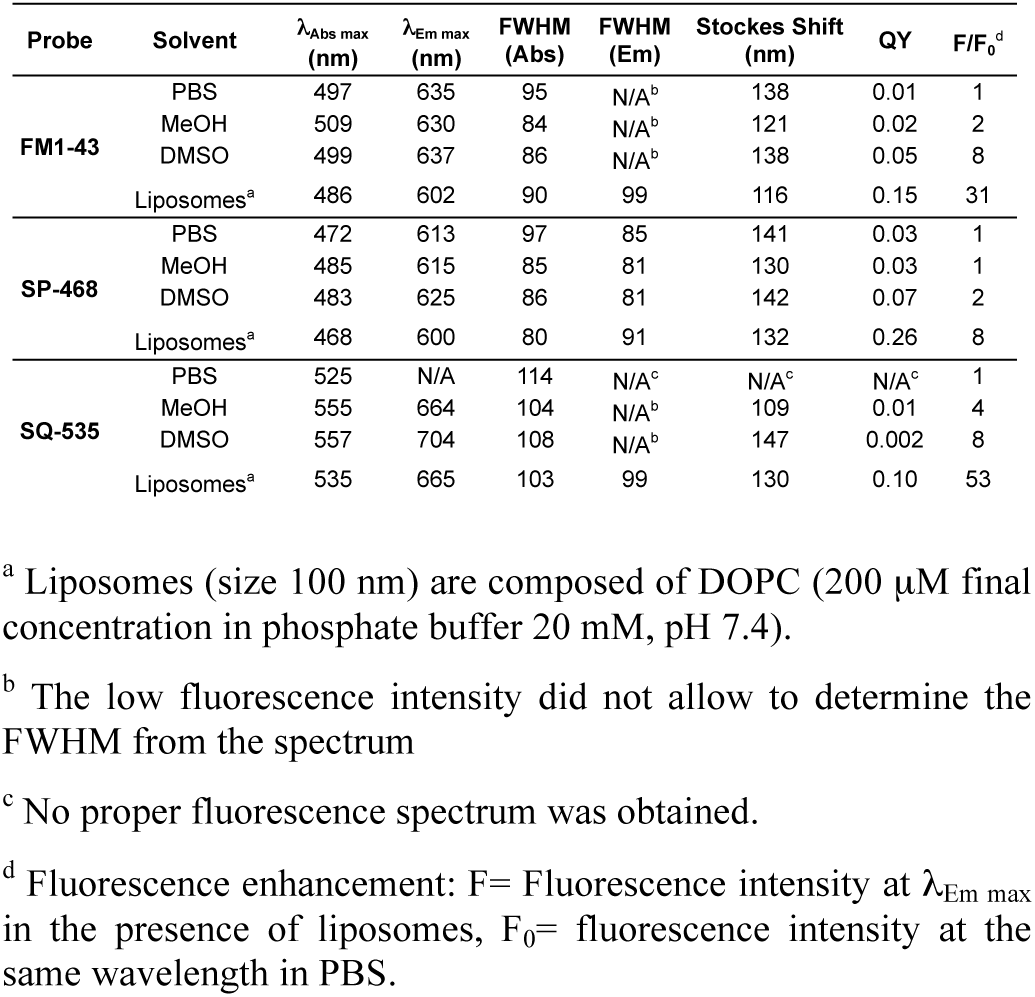
Photophysical properties of the probes.

Similarly to the parent probe FM1-43, the new probes showed hypochromism in water compared to organic solvents (Figure S1), and quantum yields in water and methanol were low (Table 1). However, the newly designed SP-468 displayed significant differences compared to FM1-43 (Figure 2A). First, a hypsochromic shift was observed in all solvents, in both absorption (up to 25 nm) and emission (up to 22 nm). As expected and discussed above, the replacement of a butyl chain for a methyl group decreased the electron density of the nitrogen donor, thus provoking the blue shift. In the presence of liposomes, the new probes showed significant blue shift in the absorption and emission spectra compared to water and polar organic solvents, similarly to that of FM1-43. However, their Stokes shift was much larger, reaching 130 nm. This larger Stokes shift could be explained by a different insertion of the probes, where the fluorophore of new probes will adopt a parallel orientation to the membrane surface.

**Figure 2.**
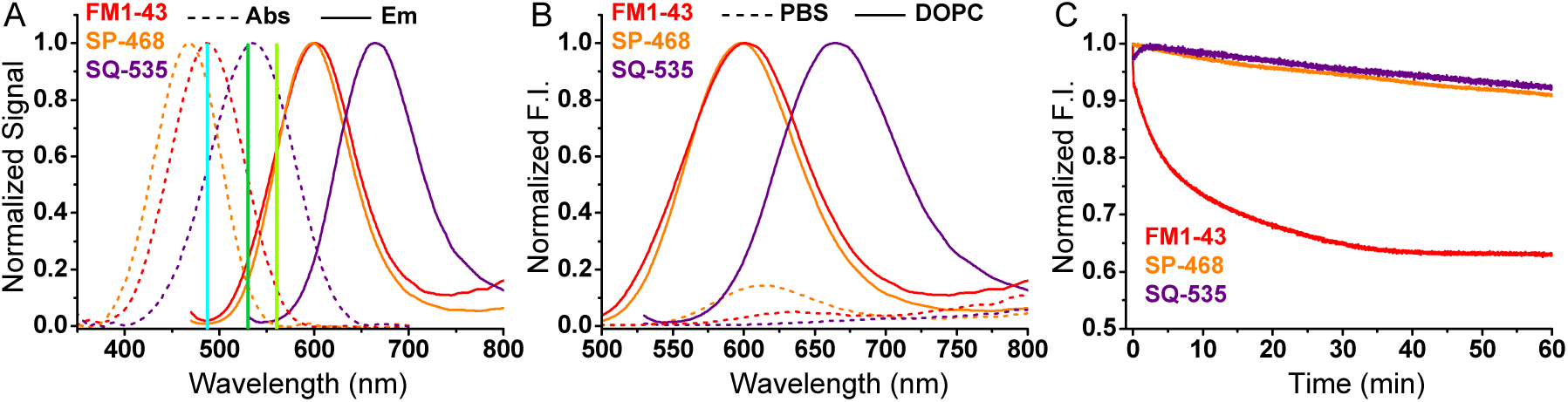
(A) Normalized absorption and emission spectra of the styryl probes in the presence of DOPC vesicles (200 μM). The 3 vertical lines indicate the most common laser lines used in imaging namely: 488, 530 and 560 nm. Absorption spectra in presence of liposomes were corrected from scattering. (B) Normalized fluorescence spectra depicting the fluorescence enhancement of the probes from PBS to insertion into DOPC vesicles (200 μM). (C) Photostability: evolution of the fluorescence intensity over the time under continuous irradiation. The probes were excited at wavelengths corresponding to the same optical density values (0.03). Concentration of the probes was 2 μM

It was noteworthy that SP-468 displayed a sharper absorption band (FWHM= 80 nm) compared to FM1-43 (90 nm) (Figure 2A), whereas both SP-468 and FM1-43 are efficiently excited by the commonly used 488-nm laser (84 and 99% respectively). Advantageously, SP-468 with a blue shifted excitation wavelength combined to a sharper peak is less prone to cross-talk compared to FM1-43. Indeed, while FM1-43 is also well excitable at 530 and 560 nm (52% and 13% respectively), SP-468 is only excited at 14 and 3% at these laser wavelengths (see laser lines on Figure 2A). The presence of liposomes resulted in significant enhancement of fluorescence intensity for all three probes, which according to the earlier studies indicates that the probe is located in highly organized environment of lipid membranes that minimize rotation-induced quenching in the styryl dyes. However, in case of SP-468, the quantum yield in PBS was slightly higher compared to FM1-43, which decreased fluorescence enhancement from PBS to liposomes (8-fold vs 31 fold, respectively, Figure 2B). The monitoring of the fluorescence signal over the time and in the presence of lipid vesicles indicated that FM1-43 and SP-468 immediately incorporated the lipid bilayer, whereas SQ-535 showed binding within less than a minute (Figure S2). We can conclude that both FM1-43 and SP-468 are present in PBS in molecular form, showing very fast binding kinetics. On the other hand, a more hydrophobic SQ-535 is probably partially aggregated in PBS, which leads to slower binding kinetics, similarly to previously reported MemBright probes.^8^ The aggregation of SQ-535 in PBS is in line with the strong blue shift in absorbance compared to organic solvents, which are much less pronounced in case of FM1-43 and SP-468. Extending the π-system from SP-468 to SQ-535 led to a 65-67 nm red shift in both absorption and emission spectra of dyes in DOPC liposomes (Figure 2A). SQ-535 constitutes a suitable PM probe with high fluorescence enhancement 53-fold, figure 2B). Although it is well excitable at 488 nm (57%), SQ-535 starts emitting at only 560 nm and thus can be used in combination with green emitting dye like BODIPY, fluorescein or eGFP (*vide infra*). We then assessed the photostability of the probes in the presence of DOPC liposomes and it was noteworthy that FM1-43 photobleached in an exponential manner (30% decrease of the fluorescence intensity after 15 min irradiation), while after 1 h SP-468 and SQ-535 lost only 10% and 8% of their fluorescence intensity respectively (Figure 2C). This difference might be assigned to the design of the probes that impose fluorophore of SP-468 and SQ-535 to locate parallel to the membrane plane, in contrast to FM1-43 where the fluorophore itself insert perpendicular to the lipid bilayer.^22^

### Cellular imaging

Prior to imaging studies, the effect of the probes on the cell viability was assessed by MTT assay. As expected, whereas SP-468 and SQ-535 did not show significant signs of cytotoxicity up to 5 μM, the cationic nature of the FM1-43 provoked a notable decrease of the cell viability after 24h at various concentrations (Figure S3). As a first imaging experiment, the ability of FM1-43 and SP-468 to stain the PM of live cells was evaluated on 4 different eukaryotic cell lines at a concentration of 1 μM in no-wash conditions. Since FM1-43 and SP-468 possess similar photophysical properties, the same image acquisition and processing settings were applied for both. The results showed that in the same conditions, whereas FM1-43 failed at providing any signal, SP-468 immediately displayed a selective and intense staining of the PM regardless the cell line (Figure 3). When the laser power was increased, the images obtained with FM1-43 were of poor quality with a weak signal/background ratio even after 20 minutes whereas SP-468 maintained a high quality staining over that same period (Figure S4). SQ-535, when excited at 560 nm, showed similar good PM staining properties (Figure S5) and thus constitutes a red shifted alternative to SP-468. KB cells stained with SP-468 and SQ-535 were then imaged under continuous laser scanning illumination for 10 minutes with negligible loss of intensity and with conserved PM staining thus demonstrating their photostability (movie 1 and 2). The PM staining efficiency of SP-468 was then assessed in fixed cells. We established that cell fixation followed by addition of the probe was preferable over the other way around and provided excellent results with no sign of internalization (Figure S6).

**Figure 3.**
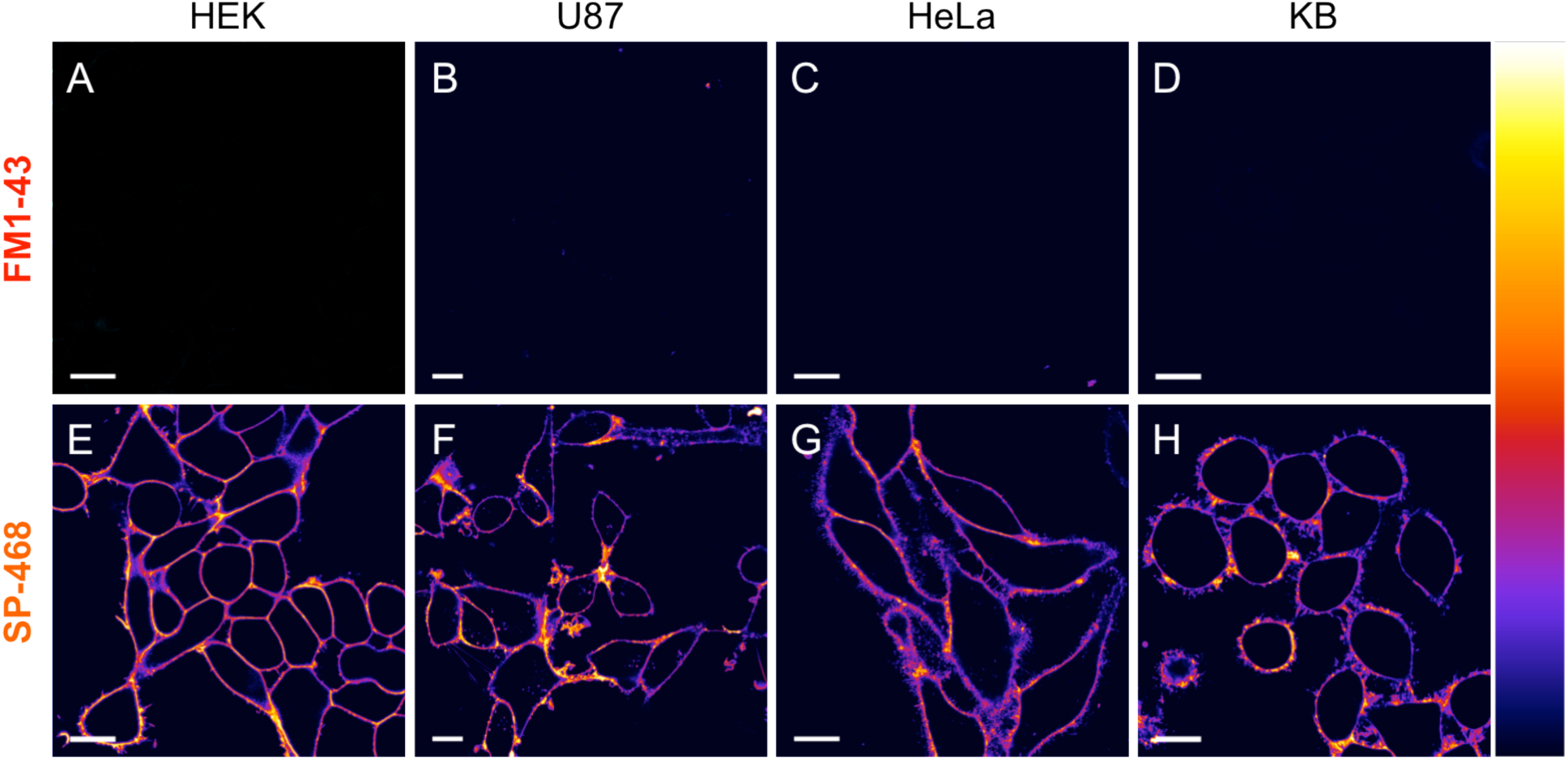
Laser scanning confocal imaging of live cells from four different cell lines stained with 1 μM of FM1-43 (A, B, C, D) and SP-468 (E, F, G, H). Images were acquired 5 minutes after addition of the dyes in the medium. Excitation wavelength was 488 nm and the fluorescence signal was collected between 520 and 700 nm. Scale bar is 15 μm. On the right is displayed the color lookup table.

Additionally, the use of SQ-535 at only 500 nM on fixed primary culture of hippocampal neurons and astrocytes in no wash conditions led to a homogeneous and bright staining of their PM (Figure 4A and B).

**Figure 4.**
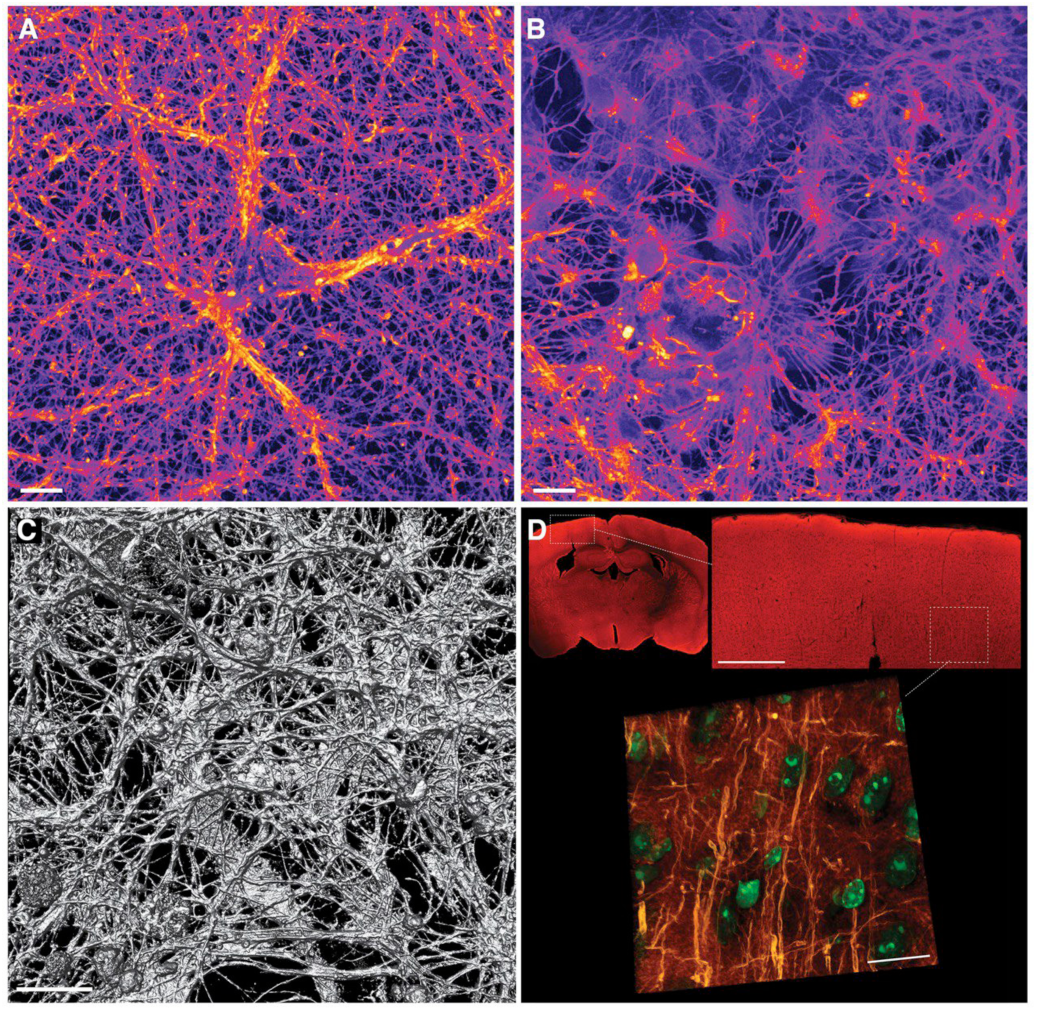
Laser scanning confocal imaging (with 20× objective) of fixed hippocampal neurons (A) and fixed astrocytic glial cells (B) incubated with SQ-535 (500 nM). (C) 3D rendering of the complex neuronal network (10 μm thick 3D volume, 93× objective). (D) Fixed brain slice images using SQ-535 (in red) and DAPI (cell nuclei in green). Some dendritic processes within the cortical layer (top right inset, scale bar 500 μm) are revealed in the 3D rendering of a 12 μm thick Z stack (scale bar: 20 μm).

The photostability and quality of staining allowed to obtaining high quality 3D images that revealed the dense neuronal network (Figure 4C). After incubation on fixed brain slices, SQ-535 provided a homogeneous staining of the whole tissue (Figure 4D). When zoomed on the cortical layer, some dendritic processes were finely revealed in a multicolor 3D rendering (Figure 4D).

### Specificity and Cellular uptake

We then aimed at comparing the specificity of the probes towards the PM. To this end, KB cells were co-stained with SQ-535 (1 mM) and FM1-43 at a higher concentration (5 mM) in order to increase its signal in imaging. After checking that crosstalk did not occur between the 2 channels in these conditions (Figure S7), a time laps imaging was performed for 10 minutes (Movie 3). The results first showed that FM1-43 at high concentration and after several minutes, provided a weak PM staining accompanied with non-specific staining of intra-cellular membranes (Figure 5A). Conversely, SQ-535 provided a fast and bright PM staining that kept its selectivity along the imaging period (Figure 5B, C and Movie 3).

**Figure 5.**
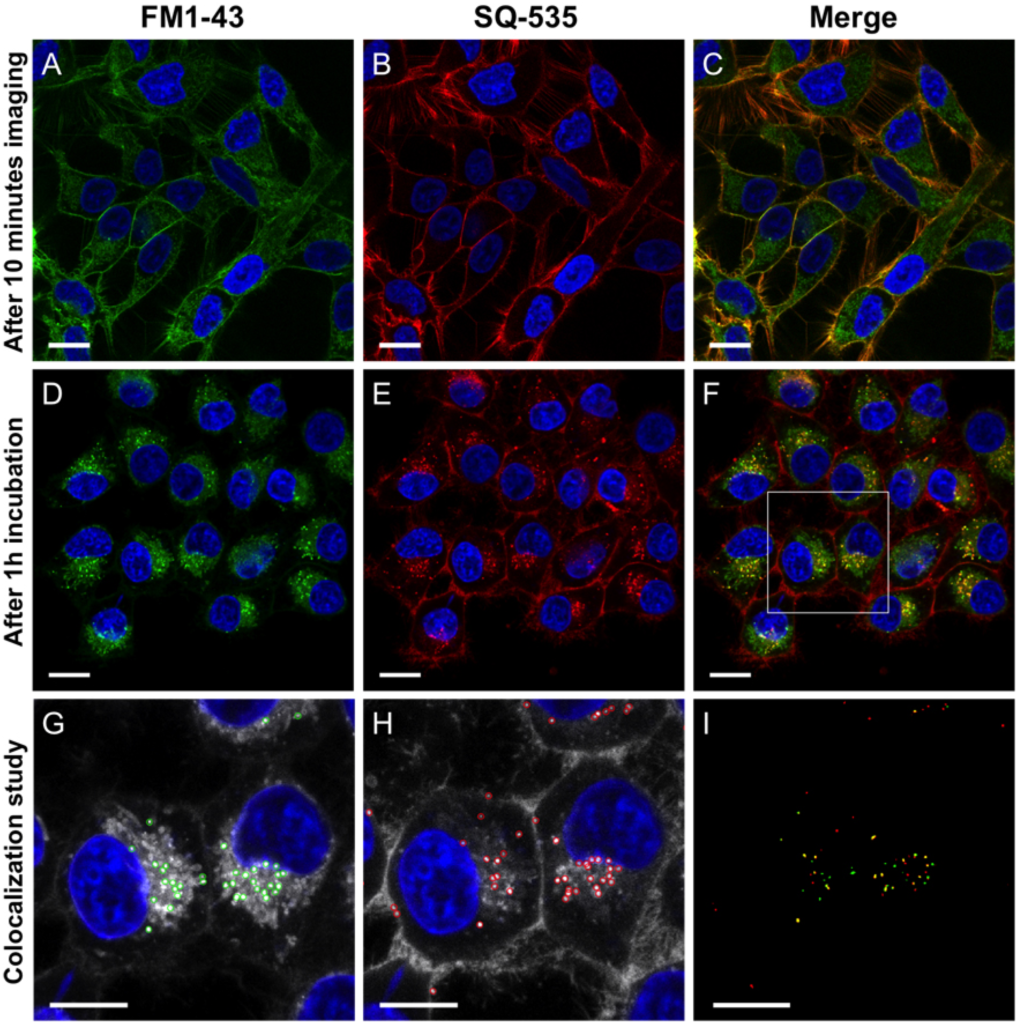
Laser scanning confocal imaging of KB cells stained with Hoechst (5 μg/mL) and 5 μM of FM1-43 (A, D, G) and 1 μM SQ-535 (B, E, H). (A, B, C) Images of live cells after 10 min of continuous imaging. (D, E, F) Images of fixed cells after 1h incubation in Opti-MEM. Merged channels (C, F) depict the different staining pattern and internalization pathway of the probes. G and H are zoomed images in the region of interest displayed in F, highlighting endosomes using spots detection plugin. (I) The resulting merged binary images of the detected endosomes depicted a weak colocalization. Scale bars are 15 μm.

In order to compare the fate of the probes over incubation, FM1-43 and SP-468 were incubated for 1h at 37°C to provoke endocytosis. Imaging revealed that FM1-43 rapidly stained intracellular membranes along with endosomes vesicles (Figure S8A, movie 4). SP-468 kept a PM staining after incubation and showed punctiform staining in the cytoplasm with characteristic movement when imaged over the time (Movie 5) depicting endosomes (Figure S8B). To assess the difference in the internalization within the same cellular experiment, FM1-43 and SQ-535 were co-incubated in KB cells for 1h at 37°C before being fixed. Analysis of images revealed that FM1-43 was actually distributed between endosomes and intracellular membranes (Figure 5D), whereas SQ-535 was distributed between the PM and endosomes (Figure 5E). Interestingly the merge images revealed that FM1-43 and SQ-535 do not fully follow the same internalization pathway as green and red signals poorly colocalized in the endosomes with a measured Pearson correlation coefficient of 0.32 (Figure 5F).

### PM staining of plant cells

FM dyes are widely used in cellular plant imaging as it efficiently stains the PM of the latter and thus is useful for segmentation and for tracking trafficking vesicles.^23, 24^ We thus investigated on the ability of our modified FM probes to conserve efficient PM staining in plant cells. For this two different experiments were conducted. On one hand, a solution of probe was injected in the leaf of 6 weeks old *Nicotiana Benthamiana* plant (Figure 6A-D); on the other hand, whole seedlings were incubated in a solution of probe to stain the roots (Figure 6 E-F). The results showed that using the same imaging and processing conditions, FM1-43 and SP-468 both stained the PM and allowed segmentation of the cells thus showing that the newly developed PM probes can be used with both mammalian and plant cells. Noteworthy, although FM1-43 and SP-468 displayed similar staining properties in the roots, FM1-43 displayed slightly more homogeneous staining in leaf cells compared to SP-468 that displayed enhanced signal in stomata (Figure 6D).

**Figure 6.**
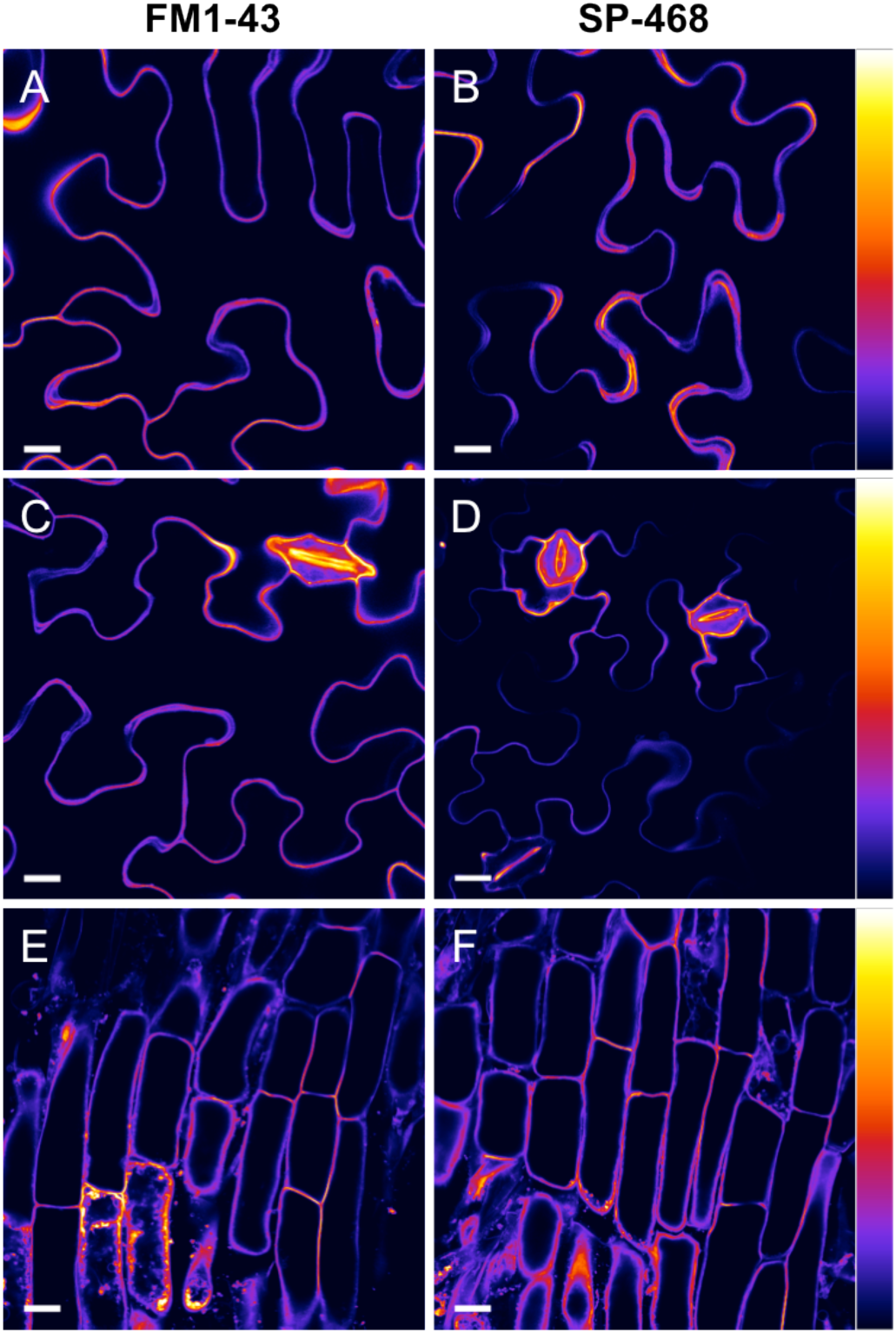
Laser scanning confocal imaging of plant cells (*Nicotiana Benthamiana*) stained with FM1-43 (A, C, E) and SP-468 (B, D, F). Images at the level of leafs without stomata (A, B) and with stomata (C, D). Images at the level of roots (E, F). Scale bar is 15 μm. On the right is the color lookup table.

## Conclusion

Although FM dyes are known as plasma membrane probes, FM1-43 displayed rather weak PM staining properties in eukaryotic cells. We showed that by conserving the same core fluorophore and by means of rational chemical modifications using adapted targeting moieties and neutralizing the cationic nature of FM1-43, styryl dyes could be efficiently converted into PM probes with interesting properties for PM bioimaging. Notable improvements were obtained as the tendency for crosstalk was reduced and the brightness as well as the photostability were significantly enhanced. The newly developed probes displayed fast, efficient and selective staining of PM in 5 different mammalian cell lines, including primary neurons, in contrast to parent FM1-43 that showed weak signal and significant intracellular staining. Remarkably, the new probes showed very good PM staining of fixed cells and brain slices. Additionally the ability to stain PM of plant cells was overall conserved making these new dyes adapted for both mammalian and plant cells experiments. The present work demonstrated the importance in the design of molecular fluorescent probes to fit with the bioimaging requirements. SP-468 and SQ-535 constitute reliable PM probes for users who want to benefit from large Stokes shift of PM markers with high photostability and to work with both live and fixed samples of cells and tissues.

## Materials and methods

### Synthesis

All starting materials for synthesis were purchased from Alfa Aesar, Sigma Aldrich or TCI Europe and used as received unless stated otherwise. FM1-43 was purchased from Biotium.^17^ The solvents were not dried unless mentioned in the protocol. DOPC Liposomes of ∼100 nm were obtained following a described protocol after several rounds of extrusion.^33^ NMR spectra were recorded on a Bruker Avance III 400 MHz spectrometer. Mass spectra were obtained using an Agilent Q-TOF 6520 mass spectrometer. Synthesis of all new compounds is described in the supporting information.

### Spectroscopy

For spectroscopy studies the concentration of the probes was 2 mM and solvents were of spectroscopic grade. The water was miliQ water. Absorption and emission spectra were recorded on a Cary 400 Scan ultraviolet–visible spectrophotometer (Varian) and a FluoroMax-4 spectrofluorometer (Horiba Jobin Yvon) equipped with a thermostated cell compartment, respectively. For standard recording of fluorescence spectra, the emission was collected 10 nm after the excitation wavelength. All the spectra were corrected from wavelength-dependent response of the detector. The quantum yields were determined by comparison with fluorescein^34^ (in NaOH 0.1 M, f= 0.95) for FM1-43 and SP-468 (excitation at 460 nm), and rhodamine B^35^ (in water, f= 0.31) for SQ-535 (excitation at 520 nm), following equation (1):

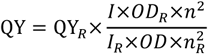

where QY is the quantum yield, I is the integrated fluorescence intensity, n is the refractive index, and OD is the optical density at the excitation wavelength. R represents the reference. The concentrations were adjusted by absorption considering the following extinction coefficients in MeOH: FM1-43, 52,000 M^-1^.cm^-1^ (according to the litterature^36^); SP-468, 50,000 M^-1^.cm^-1^ (determined from measured precursor SP-dialkyne) and SQ-535, 49,800 M^-1^.cm^-1^ (determined from measured precursor SQ-dialkyne).

### Photostability assay

The dyes (2 µM) in the presence of vesicles (100 eq of DOPC) in phosphate buffer (pH 7.4) were all excited at the same optical density value based on their absorption spectra, namely @ OD= 0.03. FM1-43 was excited at 450 nm, SP-468 at 430 nm and SQ-535 at 478 nm. Optical slits were set at 16 mm for excitation and 1 mm for emission. The emission was monitored at 600 nm for FM1-43 and SP-468 and at 660 for SQ-535.

### Membrane binding kinetics

A solution of DOPC (200 µM) vesicles in PB at pH 7.4 were stirred under excitation for 1 min, then 5 µL of probe (5 µL of stock at 400 µM in DMSO) was added. Excitation was at 460 nm for FM 1-43 and SP-468 and the fluorescence was monitored at 600 nm. For SQ-535 it was 520 nm and 660 nm respectively.

### Cellular imaging

HEK293 (ATCC® CRL-3216™) and HeLa (ATCC® CCL-2™) cells were grown in Dulbecco’s Modified Eagle Medium without phenol red (DMEM, Gibco-Invitrogen) supplemented with 10% fetal bovine serum (FBS, Lonza), 1% L-Glutamine (Sigma Aldrich) and 1% antibiotic solution (Penicillin-Streptomycin, Sigma-Aldrich) at 37°C in humidified atmosphere containing 5% CO_2_. U87 (ATCC® HTB-14™) were grown in Minimum Essential Medium (MEM, Gibco-Invitrogen) supplemented with 10% FBS, 1% Ultra-Glutamine (Gibco-Invitrogen) and 1% antibiotic solution. KB cells (ATCC® CCL-17) were grown DMEM (Gibco-Invitrogen) supplemented with 10% FBS (Lonza), 1% nonessential amino acids (Gibco-Invitrogen), 1% MEM vitamin solution (Gibco-Invitrogen), 1% L-Glutamine (Sigma Aldrich) and 0.1% antibiotic solution (gentamicin, Sigma-Aldrich) at 37°C in humidified atmosphere containing 5% CO_2_. Cells were seeded onto a chambered coverglass (IBiDi®) at a density of 5×10^4^ cells/well 24 h before the microscopy measurement. For imaging, the culture medium was removed and the attached cells were washed with Opti-MEM (Gibco– Invitrogen). Next, the cells were incubated in Opti-MEM with Hoechst (5 µg/mL) to stain the nuclei for 1h00. The cells were then washed with Opti-MEM and a freshly prepared solution of Styryl probe (2 mM in Opti-MEM) was added to the cells prior to imaging. Confocal fluorescence images of the cells were taken on a Leica TSC SPE confocal microscope using a 63 × oil immersion objective. The images were processed with Icy software.^37^ For colocalization between FM1-43 and SQ-535 the following settings were used: FM1-43 (ex: 488 nm, Em: 515-600 nm) and SQ-535 (ex: 561 nm, Em: 650-750 nm). Colocalization study was performed with Icy software: Endosomes were localized by detecting intense spots using “spot detector” plugin. The obtained binary images of endosomes were merged and the “colocalization Studio” plugin^38^ was used to determine the colocalization Pearson’s colocalization coefficient. Neuronal imaging: All animal experiments were performed in accordance with European Community guidelines legislation and reviewed by the local ethical committee of the Paris Diderot University. Hippocampal cultures from 18-day-old Sprague-Dawley rat embryos were prepared as described.^39, 40^ Cells were dissociated by treatment with 0.25% trypsin for 15 min at 37°C and plated on poly-Ornithine (1 mg/mL, Sigma) coated glass coverslips in MEM supplemented with 10% horse serum, 0.6% glucose, 0.2% sodium bicarbonate, 2 mM glutamine, and 10 IU/mL penicillin-streptomycin. After attachment neurons were grown in neurobasal medium supplemented with B27 (Thermo Fisher) conditioned on astroglia. Neurons were then imaged on Leica SP8 confocal microscope. For imaging, the medium was removed and the attached cells were washed with warm HBSS (Gibco-Invitrogen) three times. Neurons were fixed with 4% PFA, 0.1% glutaraldehyde during 10 min at 37°C. Samples were washed 3 times with HBSS and incubated with 500 nM SQ-535 and processed directly for confocal imaging in open chamber. Mice tissue: C57BL6/J mice were maintained on a 12 h light-dark cycle with ad libitum access to food and water. All animal work was conducted following protocol approved by ethical committee. Adult C57BL6 mice were euthanized and fixed with 4% paraformaldehyde by transcardial perfusion, post-fixed for 30 min in the same fixative, rinsed in PBS and kept in PBS-30% sucrose at 4°C until use. Brains were sectioned on a cryotome at 40 mm and floating sections were processed for labeling as described for cultured neurons with slight modifications: slices were washed 3 times in PBS-NH_4_Cl 50 mM and incubated 3 h with DAPI (10 mg/mL) and SQ-535 (8 µM) at room temperature under agitation. The microscope settings were: 405 nm laser for excitation of DAPI, emission was collected between 420 and 470 nm. SQ-535 was excited with 561 nm laser line and fluorescence was collected with 650-750 nm detection range. The images were processed with LAsX and Icy software.

### Cytotoxicity assay

Cytotoxicity assay was quantified by the MTT assay (3-(4,5-dimethylthiazol-2-yl)-2,5-diphenyltetrazolium bromide). A total of 1×10^4^ KB cells/well were seeded in a 96-well plate 24 h prior to the cytotoxicity assay in Dulbecco’s Modified Eagle Medium (Gibco Life Technologies -DMEM) complemented with 10% fetal bovine serum, Gentamicin (100 µg/mL), L-Glutamine (2 mM), nonessential amino acids (1 mM), MEM vitamin solution (1%) and were incubated in a 5% CO_2_ incubator at 37°C. After medium removal, an amount of 100 µL DMEM containing 5 µM, 1 µM or 0.2 µM of SMCy (SMCy3, SMCy3.5, SMCy5, SMCy5.5, SMCy7 and SMCy7.5) was added to the KB cell and incubated for 3 h at 37°C (5% CO_2_). As control, for each 96-well plate, the cells were incubated with DMEM containing the same percentage of DMSO (0,5% v/v) as the solution with the tested dyes or with Triton 1% as a positive control of cytotoxicity. After 24h of dye incubation, the medium was replaced by 100 µL of a mix containing DMEM + MTT solution (diluted in PBS beforehand) and the cells were incubated for 4 h at 37°C. Then, 75 µL of the mix was replaced by 50 µL of DMSO (100%) and gently shaken for 15 min at room temperature in order to dissolve the insoluble purple formazan reduced in living cells. The absorbance at 540 nm was measured (absorbances of the dyes at 540 nm were taken into account). Each concentration of dye was tested in sextuplicate in 3 independent assays. For each concentration, we calculated the percentage of cell viability in reference of the control DMEM+ 0.5% DMSO.

### Plant imaging

*Nicotiana Benthamiana* were used as plant model and were obtained from the Institut de biologie moléculaire des plantes (IBMP) greenhouse. Staining of leaf cells: leaves from 5-6 weeks old plants were infiltrated with the help of a syringe containing a solution of probes (5 mM in water). After 30 min, disk of leafs were taken off using a hole punch of 0.7 cm in diameter. The leaf samples were maintained in water between cover slips and were directly imaged. Staining of root cells: young plants (3 to 5 days after germination) were placed in 1 mL solution of probe (5 mM in water) for 1 hour at RT. The roots were washed with water and placed under cover slips before being imaged.

## Acknowledgements

This work was supported by the European Research Council ERC Consolidator grant BrightSens 648528 and ANR Bright-RiboProbes (ANR-16-CE11-0010). The authors would like to thank Neurimag (Paris) and Romain Vauchelles for his assistance at the PIQ platform as well as the PACSI analysis platform. We are grateful to Leducq foundation for its support concerning NeurImag SP8 confocal system purchase. We are also grateful to IBMP’s gardener and Elodie Klein for providing the plants and Dmytro Danylchuk for the preparation of liposomes.

## ASSOCIATED CONTENT

Data on synthesis protocol, ^1^H and ^13^C NMR and mass spectra, spectroscopy, additional cellular imaging, and cytotoxicity assays.

Movie 1. Photostability SP-468 (AVI)

Movie 2. Photostability SQ-535 (AVI)

Movie 3. FM1-43 & SQ-535 time laps (AVI)

Movie 4. Incubation of FM-1-43 (AVI)

Movie 5. Incubation of SP-468 (AVI)

---

Table of content graphic (TOC)

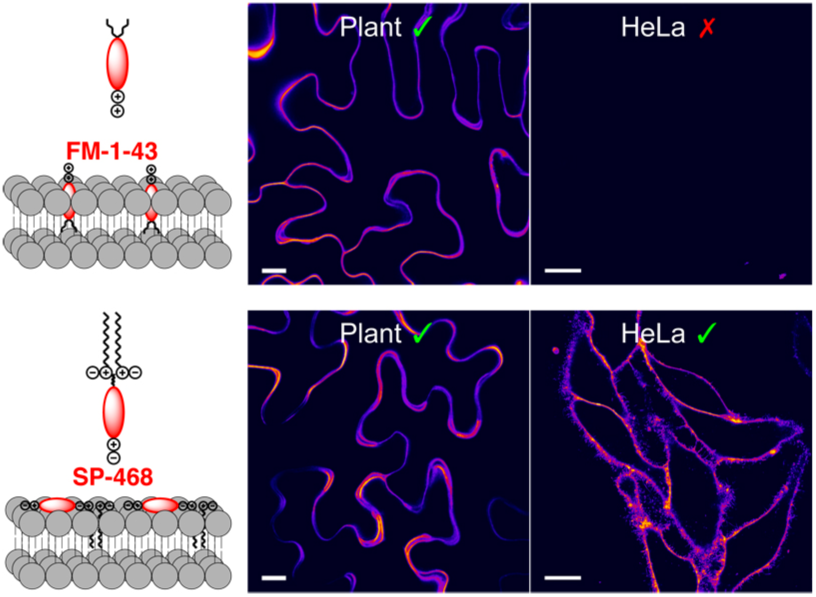

