## Supplementary information for "Molecular Tuning of Styryl Dyes Leads to Versatile and Efficient Plasma Membrane Probes for Cell and Tissue Imaging"

### Chemical synthesis

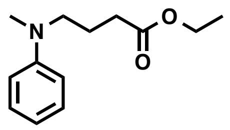

**1.** To a solution of N-methylaniline (1.00 g, 9.347 mmol) and Ethyl 4-bromobutyrate (2.70 g, 14.02 mmol, 1.5 eq) in DMSO (8 mL) was added sodium iodide (1.39 g, 9.347 mmol, 1 eq) and $NaHCO_3$ (2.30 g, 28.02 mmol, 3 eq). The solution was warmed at 80°C overnight, cooled down, extracted with EtOAc, washed with water and brine and dried over $MgSO_4$. The crude was purified by column chromatography on silica gel (100% DCM) to obtain 2.00 g of 1 (Yield=97%) as yellowish oil. Rf= 0.48 (100% DCM). The NMR was in accordance with the literature.[1]

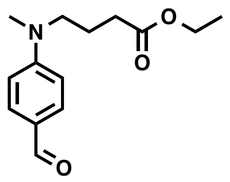

**2.** To a solution of 1 (2.00 g, 9.050 mmol) in DMF (20 mL) was slowly added at 0°C $POCl_3$ (1.6 mL). After addition the solution was allowed to warm up at room temperature and was then heated at 55°C for 2 h. The solution was slowly poured in saturated aqueous solution of $NaHCO_3$. The product was extracted with EtOAC, washed with brine and dried over $MgSO_4$. The crude was purified by column chromatography on silica gel (95/5 DCM/MeOH) to obtain 1.18 g of 2 (Yield=52%) as yellowish oil. Rf= 0.19 (100% DCM). $^1$H-NMR (400 MHz, $CDCl_3$): δ 9.75 (s, 1H, CHO), 7.75 (d, $J$ = 8.9 Hz, 2H, 2H Ar), 6.74 (d, $J$ = 8.7 Hz, 2H, 2 H Ar), 4.16 (q, $J$ = 7.1 Hz, 2H, $CH_2$ OEt), 3.49 (t, $J$ = 7.5 Hz, 2H, $CH_2$), 3.07 (s, 3H, N-$CH_3$), 2.38 (t, $J$ = 7.0 Hz, 2H, $CH_2$), 1.97 (quintet, $J$ = 7.3 Hz, 2H, $CH_2$), 1.28 (t, $J$ = 7.1 Hz, 3H, $CH_3$ OEt). $^{13}$C-NMR (101 MHz, $CDCl_3$): δ 190.1(CO aldehyde), 172.8 (CO ester), 153.4 (C Ar), 132.1 (C Ar), 125.2 (C Ar), 110.9 (C Ar), 60.6 ($CH_2$ OEt), 51.4 ($CH_2$-N), 38.4 (N-$CH_3$), 31.2 ($CH_2$), 22.1 ($CH_2$), 14.2 ($CH_3$ OEt). HRMS (ESI+), calcd for $C_{14}H_{20}NO_3$ $[M+H]^+$ 250.1443, found 250.1435.

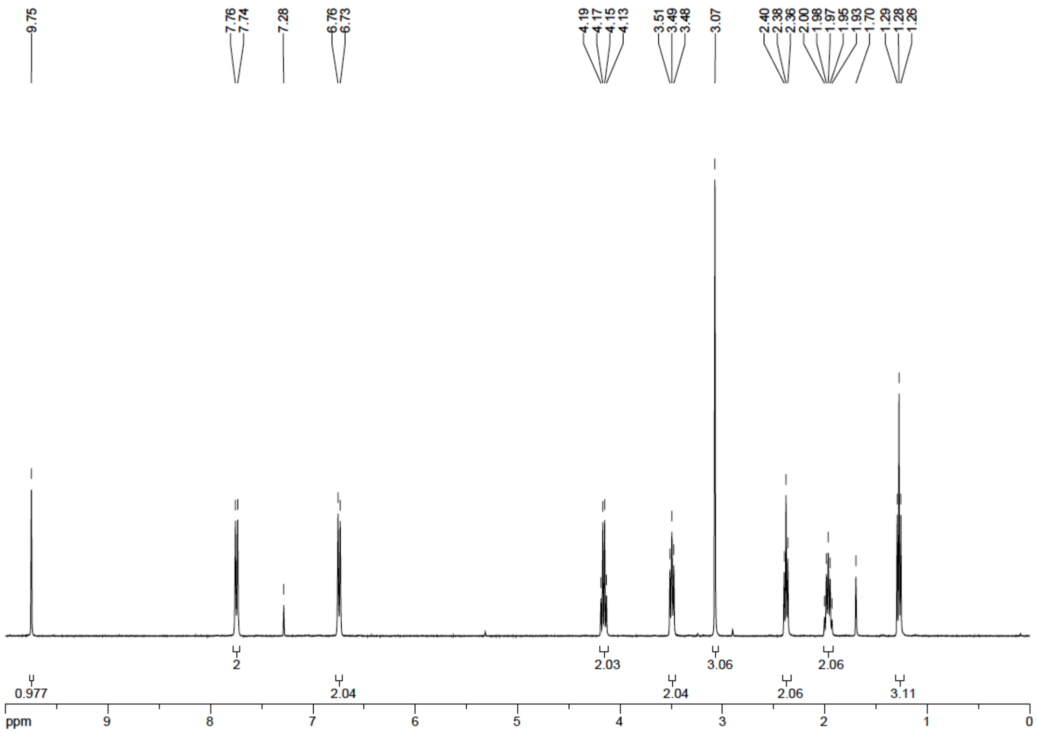


$^{1}H$ NMR spectrum of 2

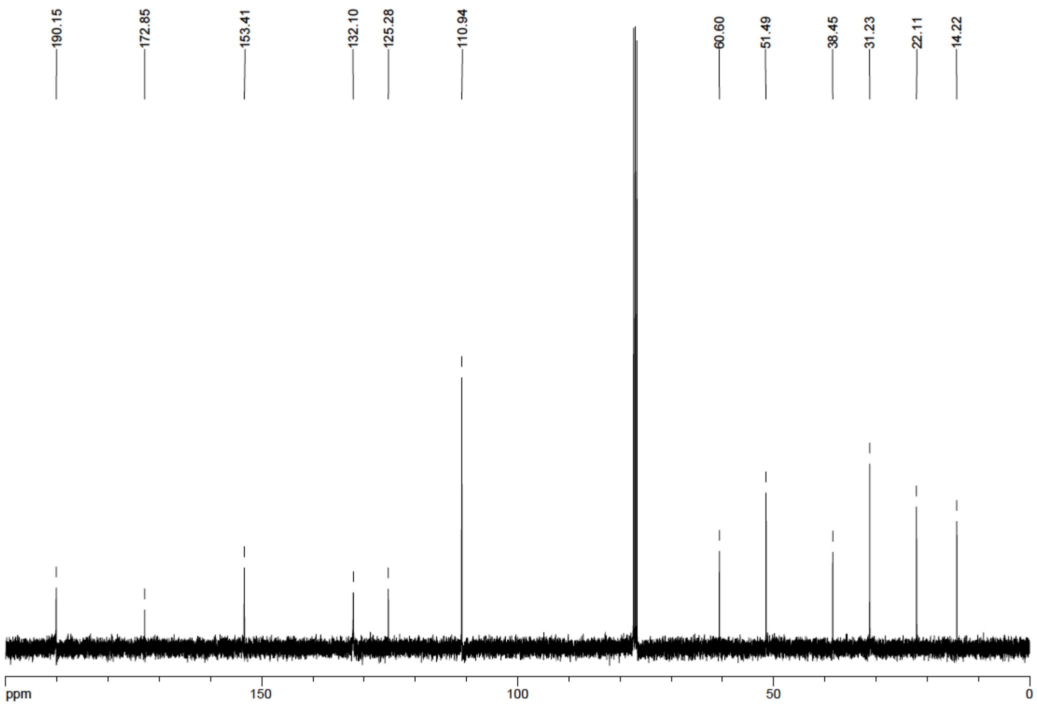


$^{13}C$ NMR spectrum of 2

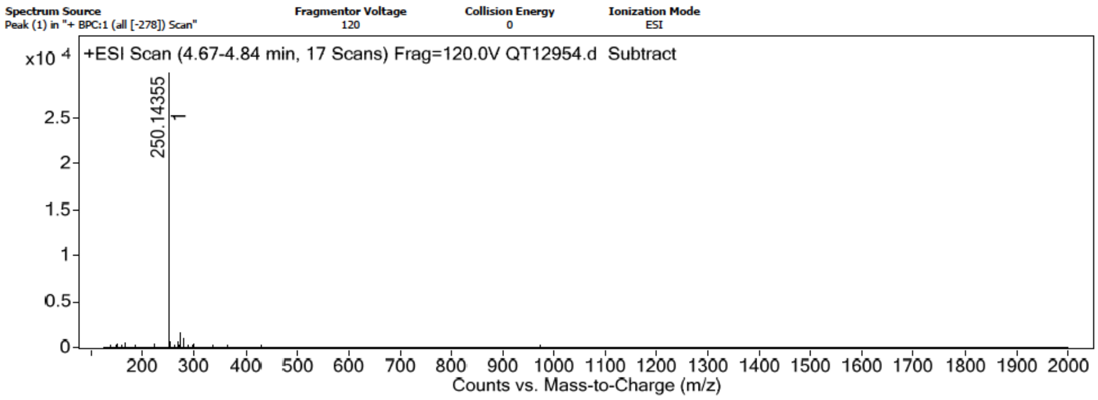


HRMS spectrum of 2

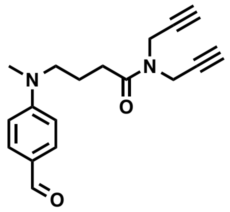

**3.** To a solution of 2 (590 mg, 2.360 mmol) in MeOH (6 mL) was added 1 mL of NaOH (5M). The solution was allowed to stir at room temperature overnight. HCl (1M) was added to the solution until the pH reach 1. The product was extracted with DCM, washed with brine and dried over $MgSO_4$. The solvents were evaporated to obtain 330 mg (1.493 mmol, 63% yield) of an orange solid. The solid was dissolved in DMF (10 mL). To this solution was added dipropargylamine (277 mg, 2.980 mmol, 2 eq), HATU (680 mg, 1.791 mmol, 1.2 eq) and DIEA (0.8 mL, 4.470 mmol, 3 eq). The solution was allowed to stir at room temperature overnight and the solvents were evaporated. The product was extracted with DCM, washed with HCl (1M) and by a saturated aqueous solution of $NaHCO_3$ before being dried over $MgSO_4$. The crude was purified by column chromatography on silica gel (95/5 DCM/EtOAc) to obtain 390 mg of 3 (Yield=56% over 2 steps) as yellowish oil. Rf= 0.31 (95/5 DCM/EtOAc). $^1$H-NMR (500 MHz, $CDCl_3$): δ 9.73 (s, 1H, CHO), 7.73 (d, $J$ = 9.0 Hz, 2H, 2 H Ar), 6.76 (d, $J$ = 9.0 Hz, 2H, 2 H Ar), 4.35 (d, $J$ = 2.3 Hz, 2H, N-$CH_2$-C≡C), 4.16 (d, $J$ = 2.2 Hz, 2H, N-$CH_2$-C≡C), 3.52 (t, $J$ = 7.5 Hz, 2H, N-$CH_2$), 3.07 (s, 3H, N-$CH_3$), 2.47 (t, $J$ = 6.7 Hz, 2H, $CH_2$-CO), 2.30 (dt, $J$ = 2.3 Hz, 2H), 2.27 (dt, $J$ = 2.3 Hz, 2H), 2.01 (m, 2H, $CH_2$). $^{13}$C-NMR (126 MHz, $CDCl_3$): δ 190.15 (CO aldehyde), 171.32(CO amide), 153.51 (C Ar), 132.13 (Cq Ar), 125.19 (Cq Ar), 111.02 (C Ar), 78.30 (C≡CH), 77.65 (C≡CH), 73.11 (C≡CH), 72.42 (C≡CH), 51.46 ($CH_3$), 38.40 (N-$CH_2$), 36.19 (N-$CH_2$C≡C), 34.20 (N-$CH_2$C≡C), 29.67 ($CH_2$), 21.87 ($CH_2$). HRMS (ESI+), calcd for $C_{18}H_{21}N_2O_2$ $[M+H]^+$ 297.1603, found 297.1604.

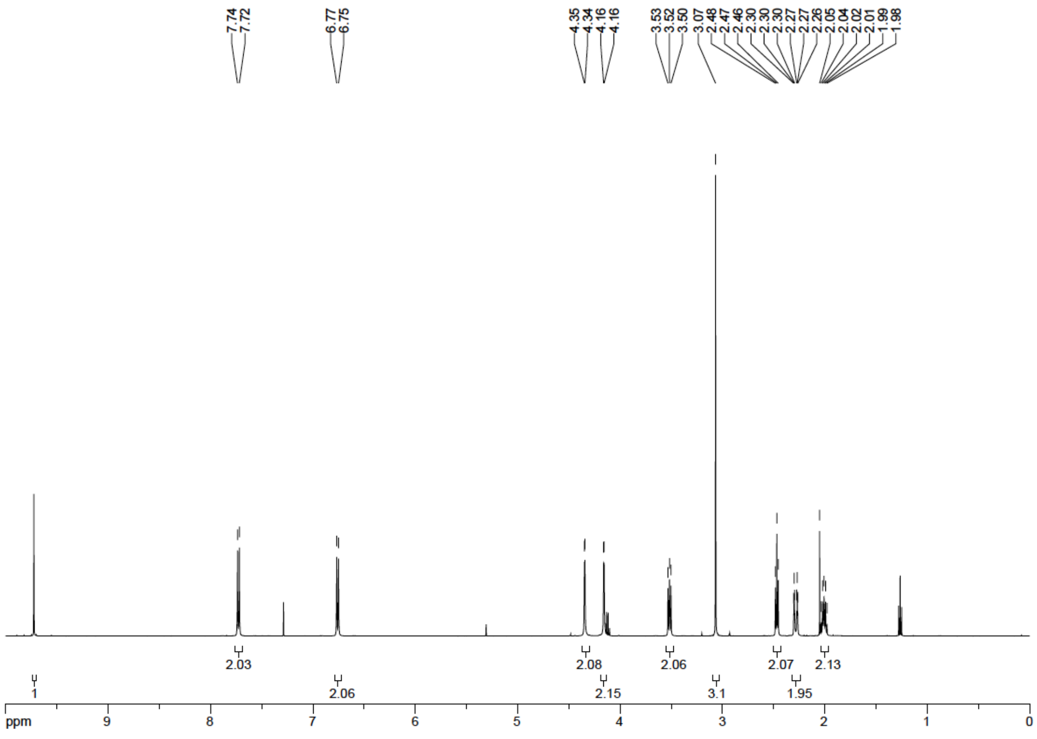


$^{1}H$ NMR spectrum of 3

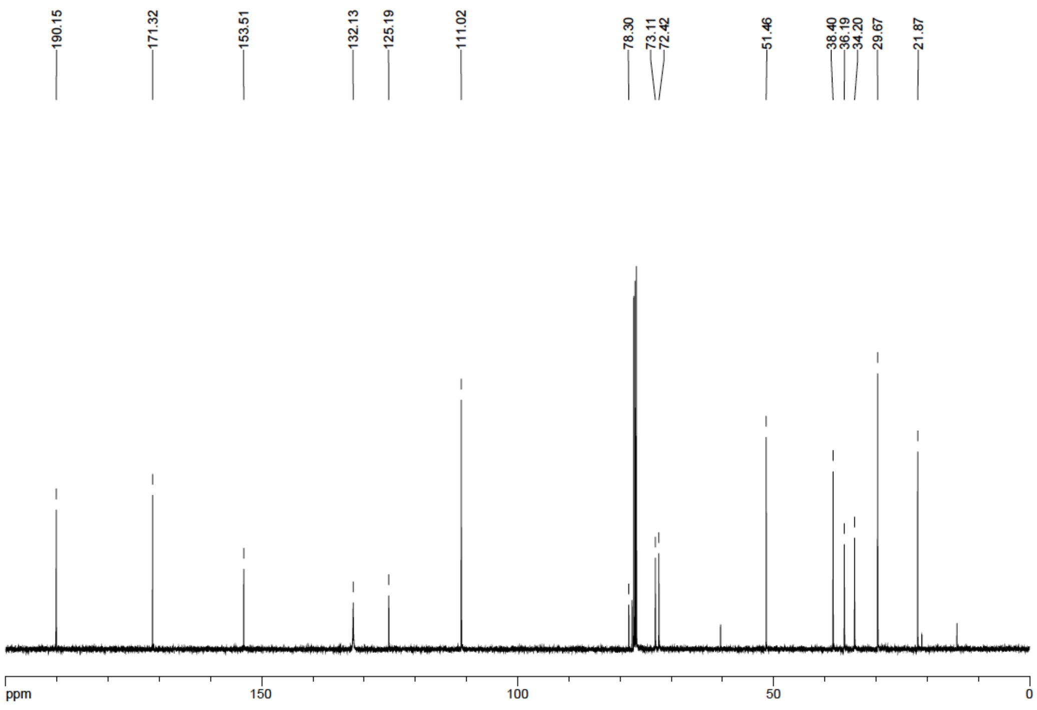


$^{13}C$ NMR spectrum of 3

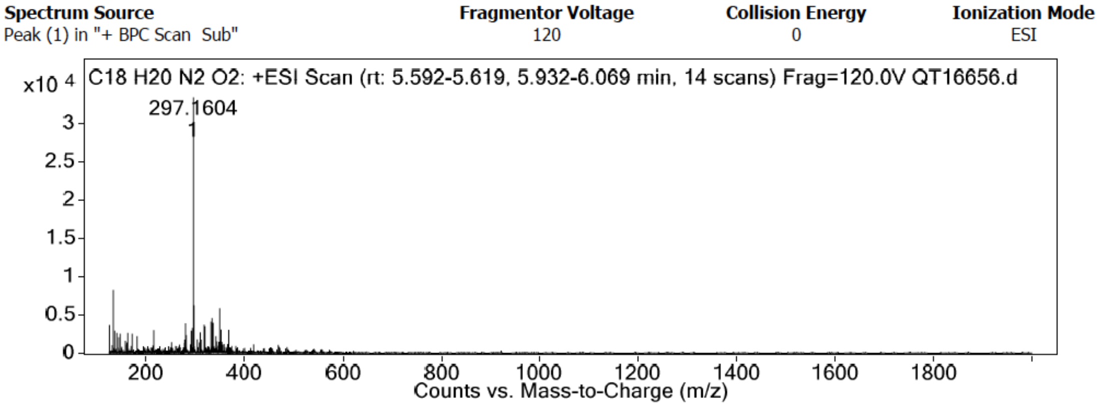


HRMS spectrum of 3

Picolinium[2] and Lepidinium[3] were synthesized according to published protocols.

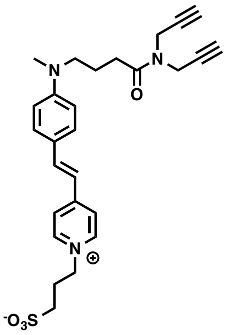

**SP-dialkyne.** To a solution of 3 (50 mg, 168.7 mmol) and picolinium (36 mg, 168.7 mmol, 1 eq) in MeOH was added 3 drops of piperidin and the solution was allowed to stir at 110°C for 4 h. The solvents were evaporated and the crude was purified by column chromatography on silica gel (95/5 to 85/15 DCM/MeOH) to obtain 21 mg of SP-dialkyne (Yield=25%) as orange amorphous solid. Rf= 0.11 (9/1 DCM/MeOH). $^{1}$H-NMR (500 MHz, MeOD): δ 8.60 (d, *J* = 7.0 Hz, 2H, 2 H Py), 7.96 (d, *J* = 7.0 Hz, 2H, 2 H Py), 7.81 (d, *J* = 16.0 Hz, 1H, HC=C), 7.60 (d, *J* = 9.0 Hz, 2H, 2 H aniline), 7.06 (d, *J* = 16.0 Hz, 1H, HC=C), 6.83 (d, *J* = 9.0 Hz, 2H, 2H, 2 H aniline), 4.63 (t, *J* = 7.3 Hz, 2H, $CH_2N^+$), 4.31 (dd, *J* = 9.4, 2.3 Hz, 4H N-$CH_2$-C≡C), 3.51 (t, *J* = 7.6 Hz, 2H, N-$CH_2$), 3.07 (s, 3H, N-$CH_3$), 2.88 (t, *J* = 6.9 Hz, 2H, $CH_2$), 2.81 (t, *J* = 2.3 Hz, 1H, C≡CH), 2.71 (t, *J* = 2.4 Hz, 1H, C≡CH), 2.56 (t, *J* = 6.9 Hz, 2H, $CH_2$), 2.42 (t, *J* = 7.2 Hz, 2H, $CH_2$), 1.96 (t, *J* = 7.5 Hz, 2H, $CH_2$). $^{13}$C-NMR (126 MHz, MeOD): δ 172.7(CO), 155.0 (Cq Ar), 151.6 (Cq Ar), 143.0 (C Ar), 143.0 (Cq Ar), 130.4 (C Ar), 122.6 (HC=C), 122.3 (C Ar), 116.2 (HC=C), 111.6 (C Ar), 77.8 (C≡CH), 77.5 (C≡CH), 73.1 (C≡CH), 72.2 (C≡CH), 58.0 ($CH_2$), 50.9 ($CH_2$), 46.8 ($CH_2$), 37.1 (N-$CH_3$), 36.0 (N-$CH_2$), 33.8 (N-$CH_2$), 29.4 ($CH_2$), 26.6 ($CH_2$), 21.7 ($CH_2$). HRMS (ESI+), calcd for $C_{27}H_{32}N_3O_4S$ $[M+H]^+$ 494.2114, found 494.2120.

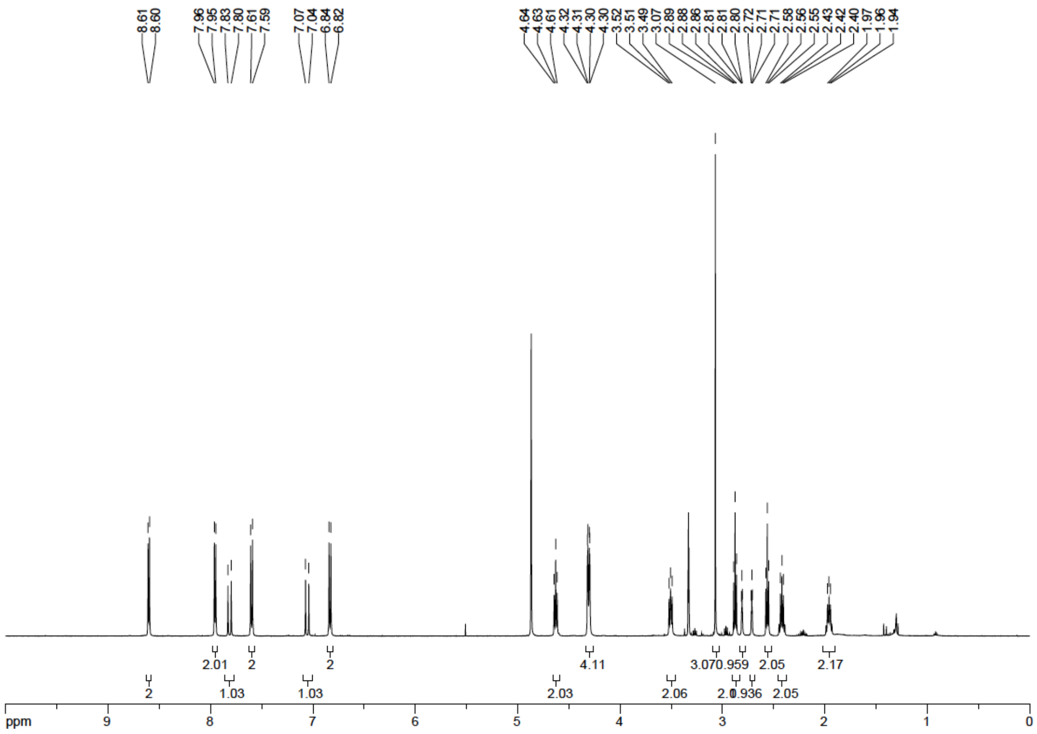


$^{1}$H NMR spectrum of SP-dialkyne

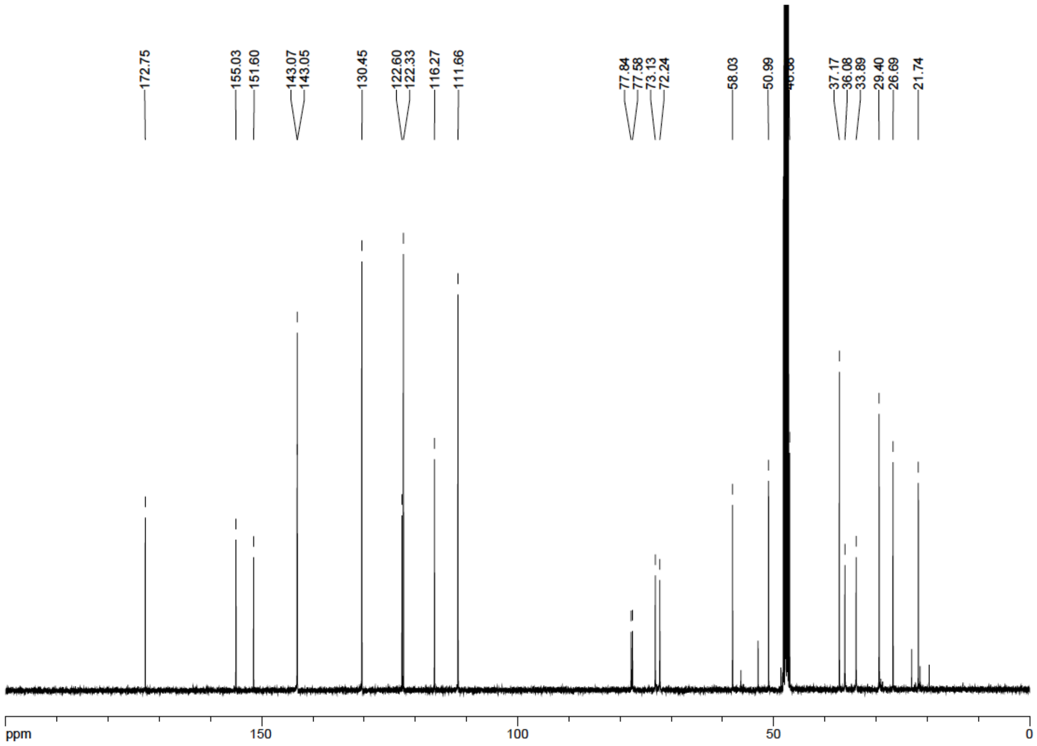


$^{13}$C NMR spectrum of SP-dialkyne

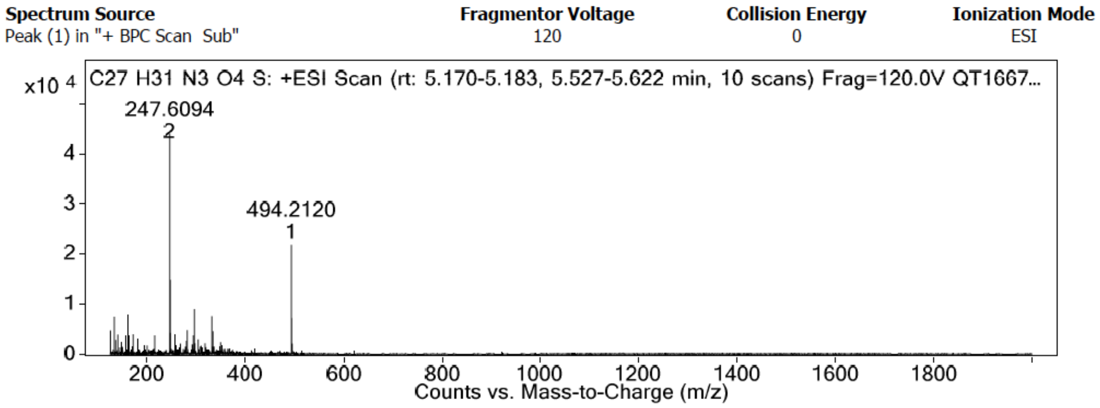


HRMS spectrum of SP-dialkyne

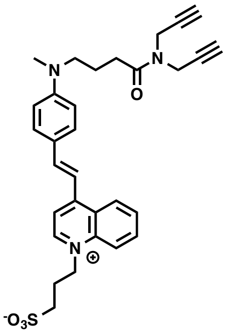

**SQ-dialkye.** To a solution of 3 (50 mg, 168.7 μmol) and lepidinium (36 mg, 168.7 μmol, 1 eq) in MeOH was added 3 drops of piperidin and the solution was allowed to stir at 110°C for 4 h. The solvents were evaporated and the crude was purified by column chromatography on silica gel (95/5 to 85/15 DCM/MeOH) to obtain 48 mg of SQ-dialkye (Yield=52%) as deep violet amorphous solid. Rf= 0.32 (9/1 DCM/MeOH). ε(MeOH)= 49,800 $M^{-1}.cm^{-1}$. $^{1}$H-NMR (500 MHz, MeOD): δ 8.84 (d, *J* = 6.7 Hz, 1H, H Lepidinium), 8.67 (d, *J* = 8.4 Hz, 1H, H Lep), 8.37 (d, *J* = 8.9 Hz, 1H, H Lep), 8.10-8.04 (m, 2H, 2H Lep), 7.87-7.81 (m, 2H, 1 H Ar, 1 HC=C), 7.67 (d, *J* = 15.5 Hz, 1H Lep, 1 HC=C), 7.58 (d, *J* = 8.8 Hz, 2H, 2 H Ar aniline), 6.76 (d, *J* = 8.9 Hz, 2H, 2 H Ar aniline), 4.99 (t, *J* = 7.8 Hz, 2H, $CH_2$-$N^+$), 4.33 (t, *J* = 3.1 Hz, 4H, N-$CH_2$-C≡C), 3.48 (t, *J* = 7.7 Hz, 2H, N-$CH_2$), 3.06 (s, 3H, N-$CH_3$), 2.99 (t, *J* = 6.7 Hz, 2H, $CH_2$), 2.84 (s, 1H, C≡CH), 2.73 (s, 1H, C≡CH), 2.58 (t, *J* = 6.8 Hz, 2H, $CH_2$), 2.45 (quintet, *J* = 7.4 Hz, 2H, $CH_2$), 1.95 (quintet, *J* = 7.4 Hz, 2H, $CH_2$). $^{13}$C-NMR (126 MHz, MeOD): δ 172.6 (CO), 154.1 (C Ar), 151.9 (C Ar), 145.3 (C Ar), 145.0 (C Ar), 138.1 (C Ar), 134.7 (C Ar), 131.2 (C Ar), 128.3 (C Ar), 126.4 (C Ar), 125.9 (C Ar), 123.0 (C Ar), 118.2 (C Ar), 113.6 (C Ar), 112.2 (C Ar), 111.7 (C Ar), 77.8 (C≡CH), 77.6 (C≡CH), 73.1 (C≡CH), 72.2 (C≡CH), 54.7 ($CH_2$), 51.1 ($CH_2$), 37. (N-$CH_3$), 36.1 (N-$CH_2$), 33.9 (N-$CH_2$), 29.4 ($CH_2$), 25.3 ($CH_2$), 21.7 ($CH_2$). HRMS (ESI+), calcd for $C_{31}H_{34}N_3O_4S$ $[M+H]^+$ 544.2270, found 544.2242.

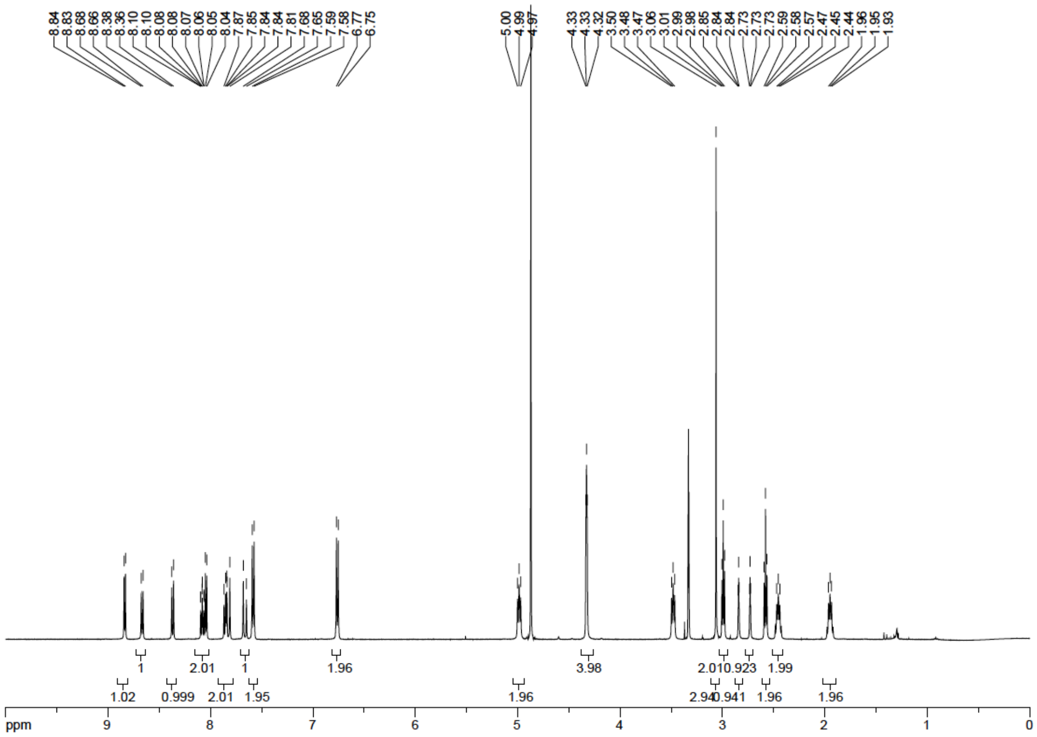


[1]H NMR spectrum of SQ-dialkyne

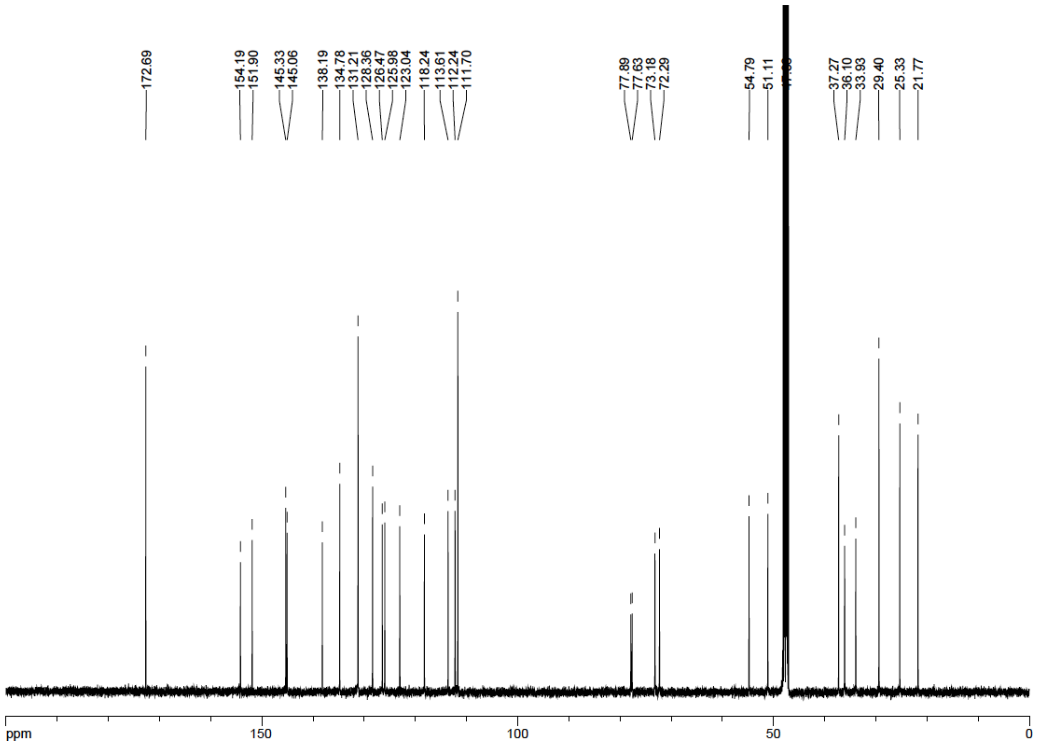


[13]C NMR spectrum of SQ-dialkyne

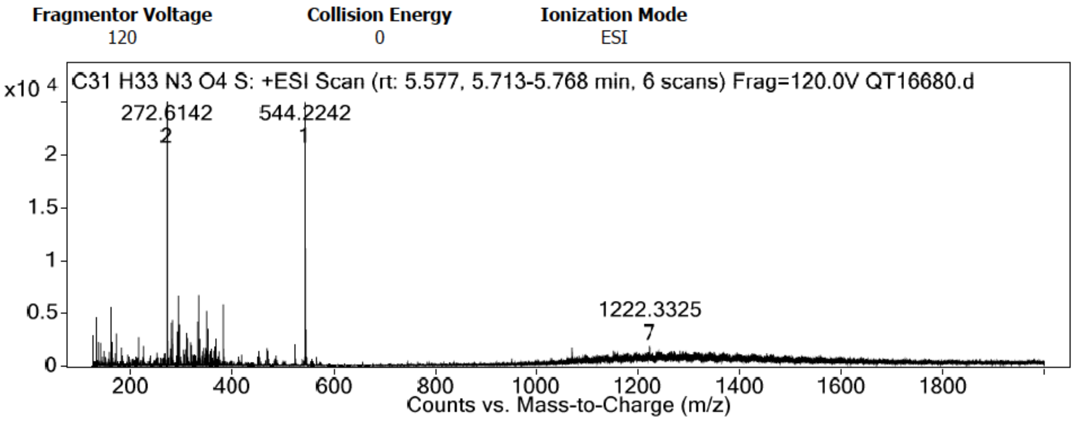


HRMS spectrum of SQ-dialkyne

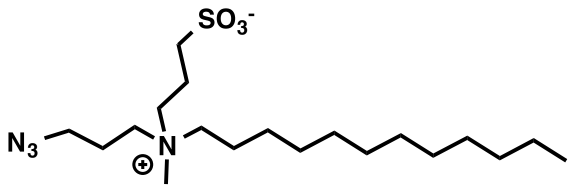


- The synthesis of CAZ was previously described[4]

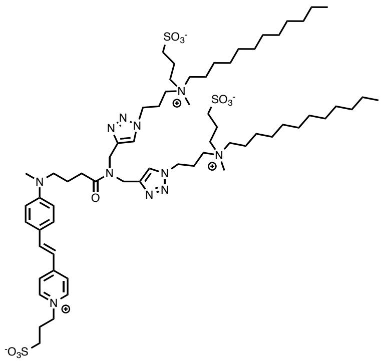

**SP-468.** To a solution of SP-dialkyne (20.0 mg, 40.52 μmol) and CAZ (40 mg, 97.24 μmol, 2.4 eq) in DMF (6 mL) was added a heterogeneous solution of $CuSO_4.5H_2O$ (10 mg, 40.52 μmol, 1 eq) and sodium ascorbate (8 mg, 40.52 μmol, 1 eq) in water (0.2 mL). The mixture was stirred at 60°C for 4 h before being evaporated. The crude product was purified by size-exclusion chromatography (DCM-MeOH 1:1) providing pure SP-468 with no sign of starting material. Due to the amphiphilic nature of the product, only $^1H$ NMR spectrum and high-resolution mass spectroscopy are provided. HRMS (ESI+), calcd for $C_{65}H_{111}N_{11}O_{10}S_3Na$ $[M+Na]^+$ 1324.7575, found 1324.7554.

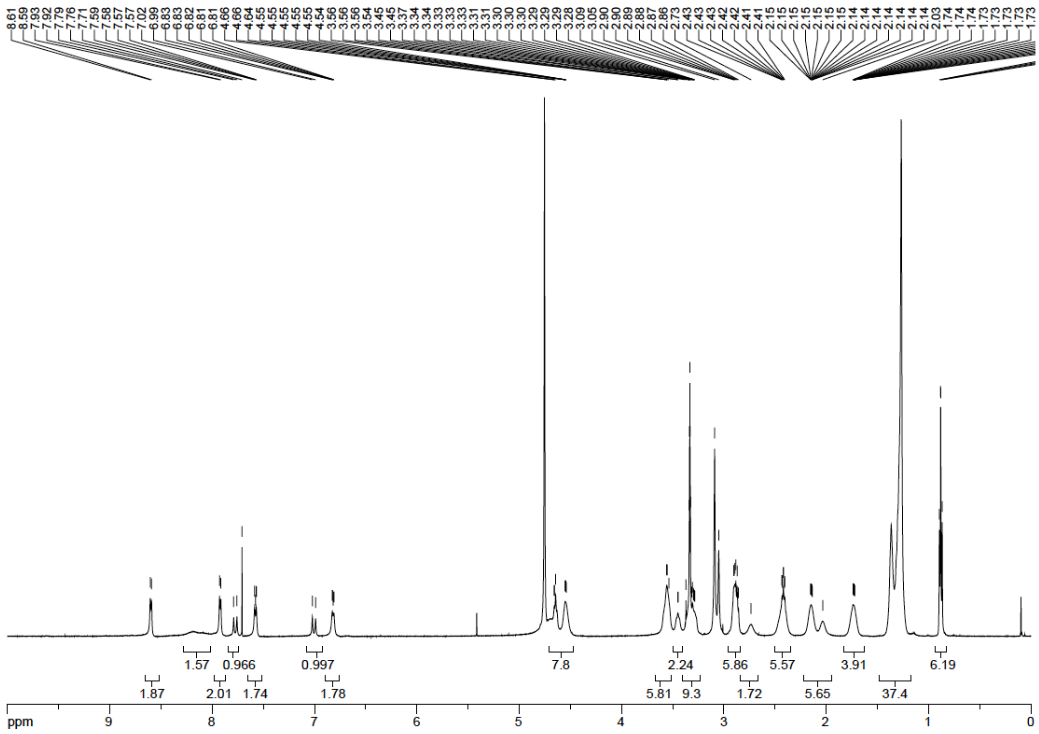

$^{1}$H NMR spectrum of SP-468

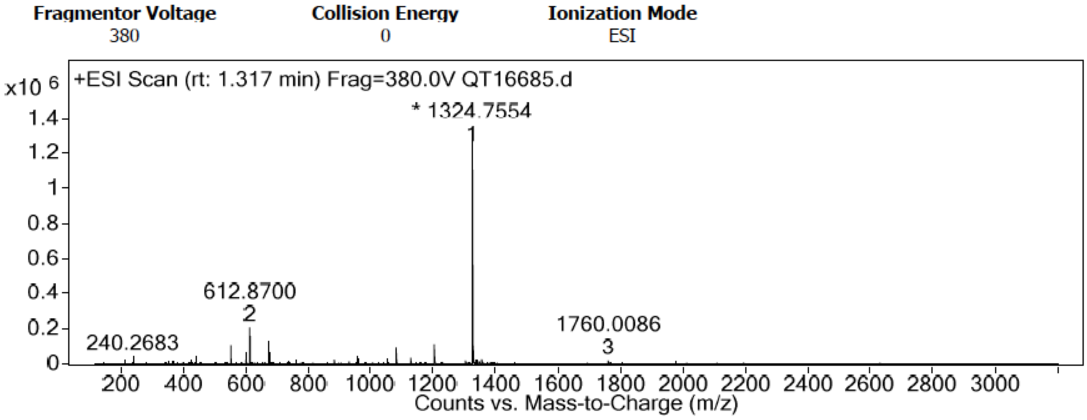


HRMS spectrum of SP-468

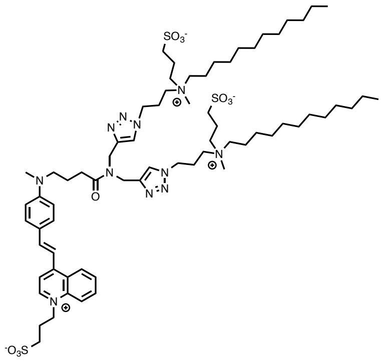

**SQ-535.** To a solution of SQ-dialkyne (20.0 mg, 36.79 μmol) and CAZ (36 mg, 88.29 μmol, 2.4 eq) in DMF (6 mL) was added a heterogeneous solution of $CuSO_4.5H_2O$ (9 mg, 36.79 μmol, 1 eq) and sodium ascorbate (7 mg, 36.79 μmol, 1 eq) in water (0.2 mL). The mixture was stirred at 60°C for 4 h before being evaporated. The crude product was purified by size-exclusion chromatography (DCM-MeOH 1:1) providing pure SQ-535 with no sign of starting material. Due to the amphiphilic nature of the product, only $^1H$ NMR spectrum and high-resolution mass spectroscopy are provided. HRMS (ESI+), calcd for $C_{69}H_{113}N_{11}O_{10}S_3Na$ $[M+Na]^+$ 1374.7732, found 1374.7665.

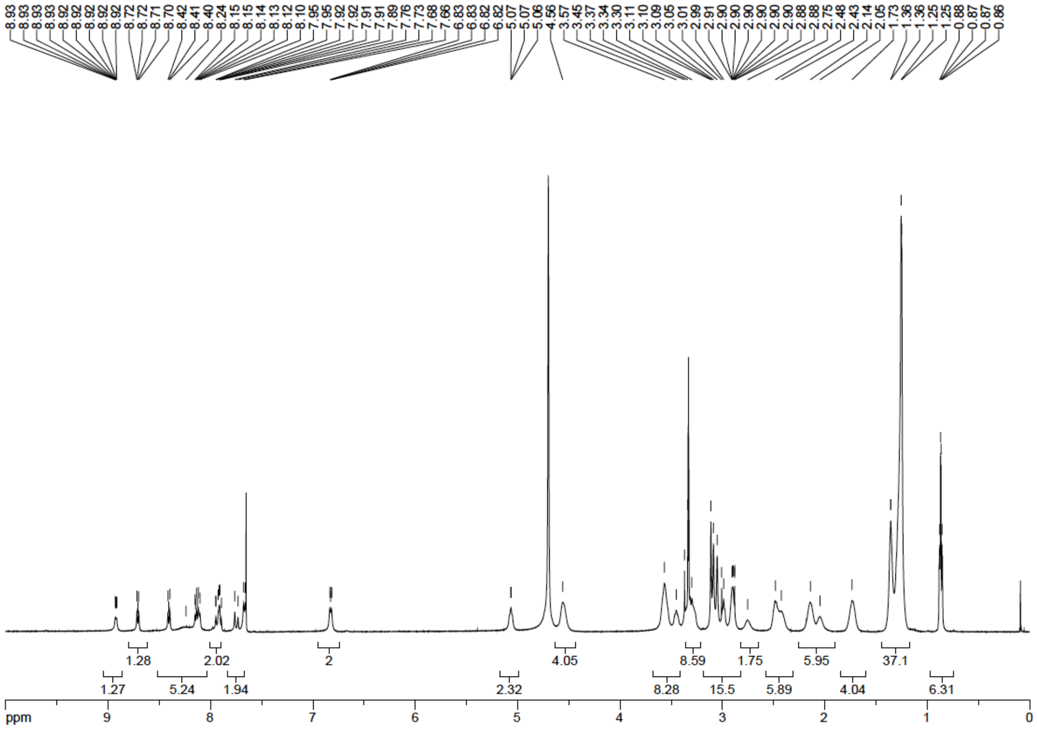


$^1H$ NMR spectrum of SQ-535

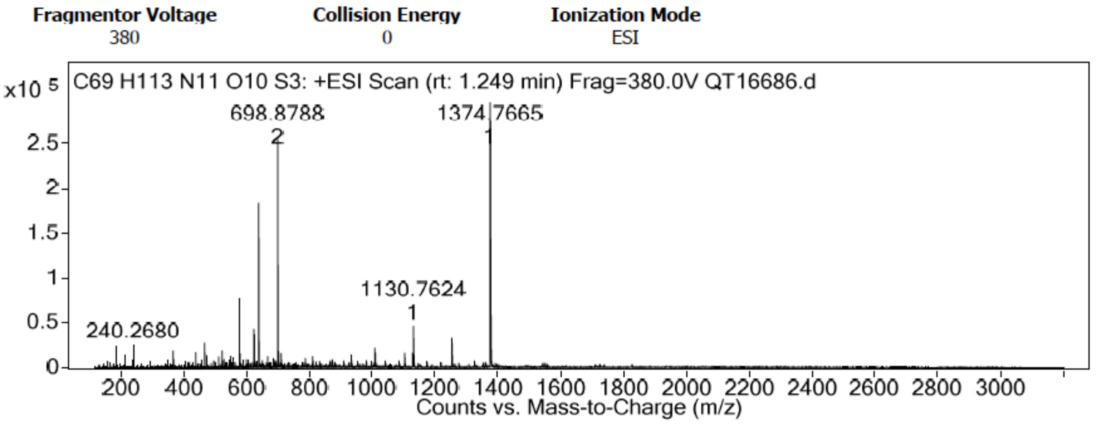


HRMS spectrum of SQ-535

### Spectroscopy

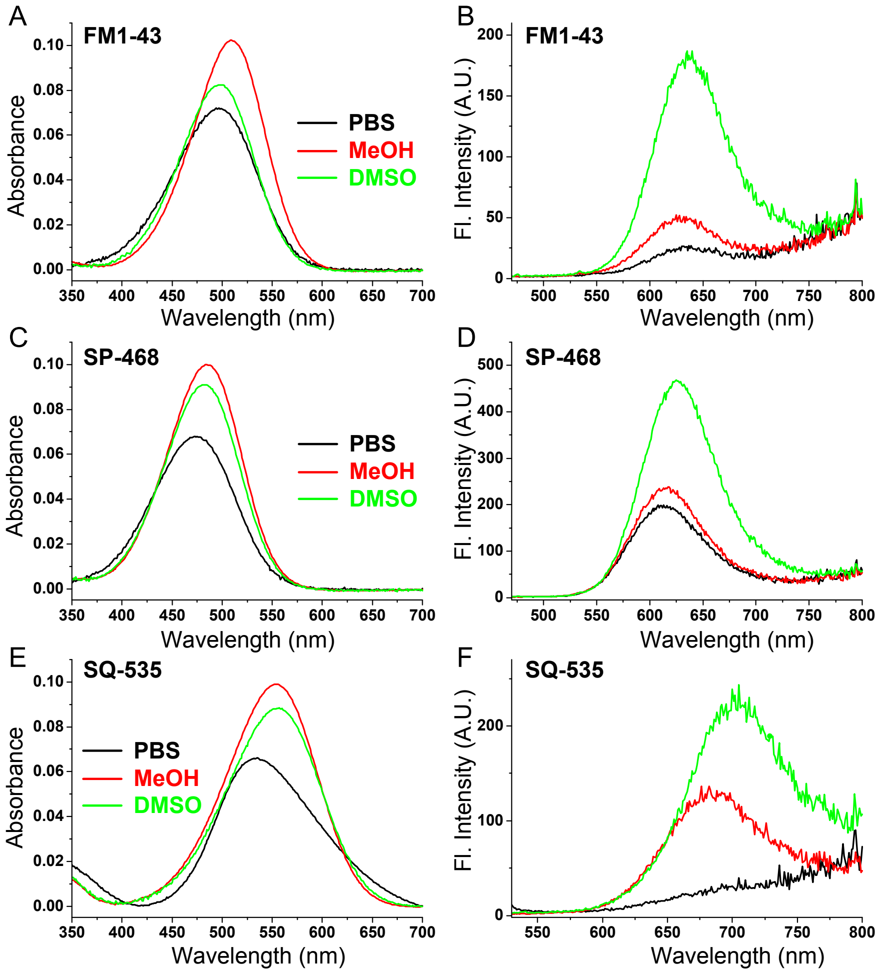


**Figure S1.** Absorption (A, C, E) and emission (B, D, F) spectra of the 3 styryl probes in various solvents. Excitation wavelength was 480 nm for FM1-43 and SP-468 and 530 nm for SQ-535. Concentration of probes was 2 μM.

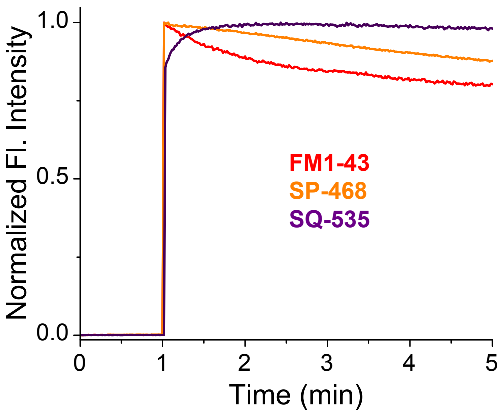


**Figure S2.** Kinetics of binding of styryl probes (2 μM) to DOPC liposomes (200 μM based on DOPC concentration). The emission was monitored over the time and at 1 min, 5 μL of probe (Stock solution DMSO at 400 μM) was added to a stirring solution of DOPC liposomes. Excitation wavelength was 480 nm for FM1-43 and SP-468 and 530 nm for SQ-535.

### Cytotoxicity

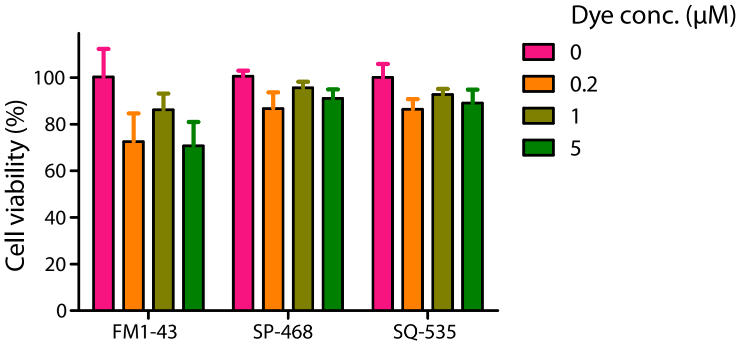


**Figure S3.** Cell viability in the presence of the styryl probes performed by MTT assay.

### Cellular imaging

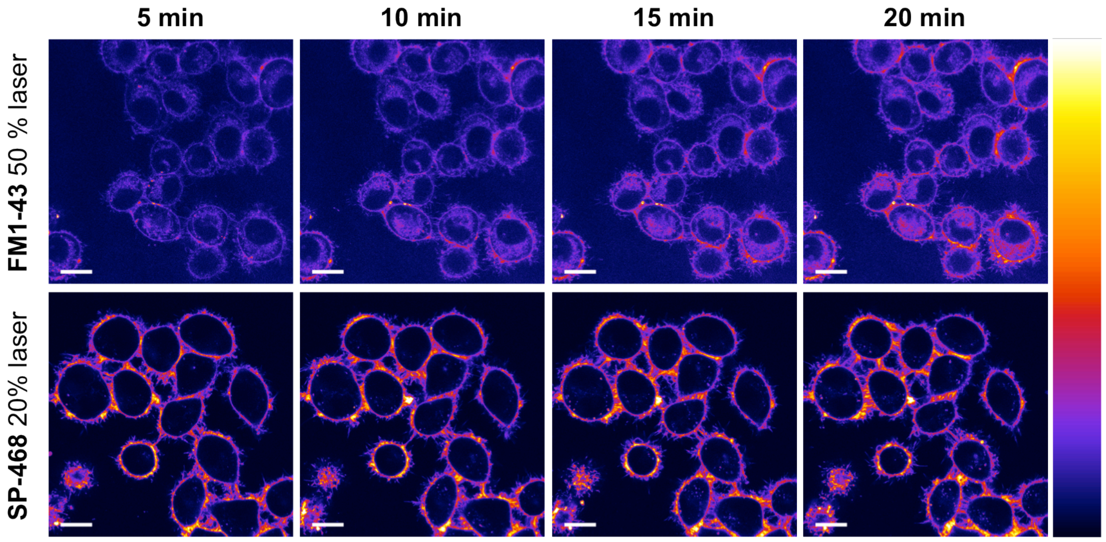


**Figure S4.** Laser scanning confocal imaging of live KB cells stained with FM1-43 and SP-468 at 1 μM. images were acquired every 5 minutes. Scale bar is 15 μm. On the right is displayed the color lookup table.

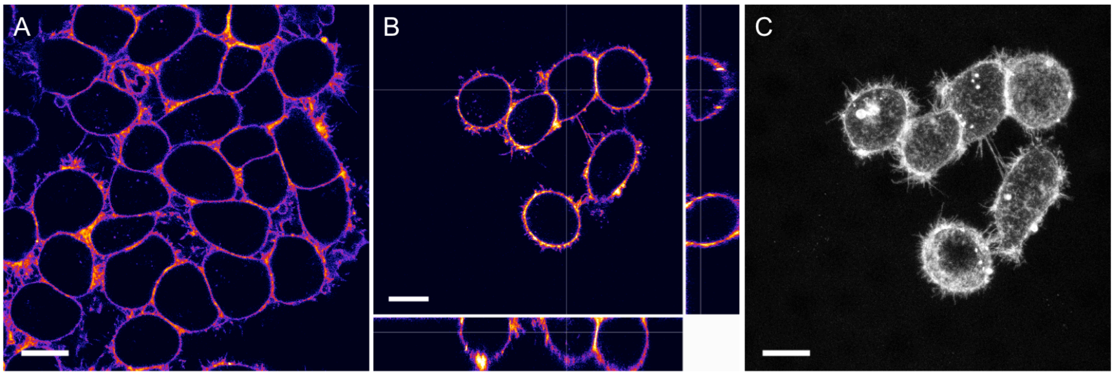


**Figure S5.** (A) Laser scanning confocal imaging of live KB cells stained with SQ-535 at 1 μM. (B) Orthogonal projection obtained by Z stack (68 frames) showing the selective staining of the PM. (C) Max projection obtained from the Z stack. images were acquired right after addition of the probe. Excitation wavelength was 560 nm and the signal was collected from 570 to 750 nm. Scale bar is 15 μm.

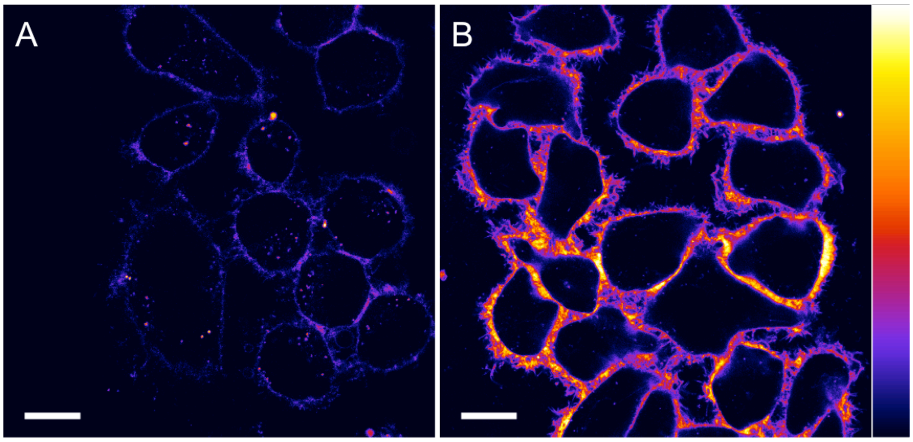


**Figure S6.** Laser scanning confocal imaging of fixed KB cells (4% PFA at room temperature for 5 minutes) with 1 μM of SP-468. (A) Staining was performed before fixation and (B) the staining was performed after fixation. Excitation wavelength was 488 nm and the fluorescence signal was collected between 520 and 700 nm. Scale bar is 15 μm. On the right is displayed the color lookup table.

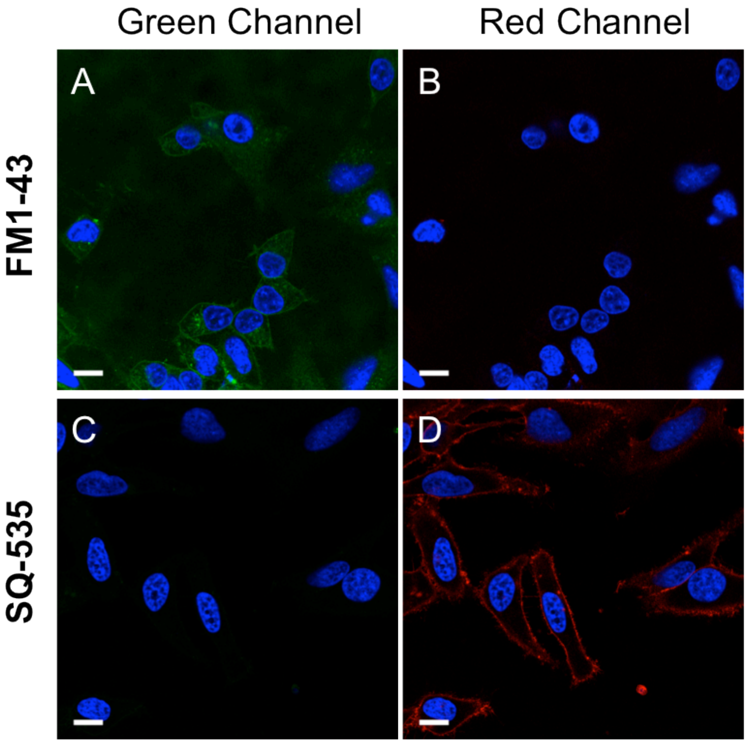


**Figure S7.** Check of cross talk between FM1-43 and SQ-535. HeLa cells separately stained with 5 μM FM1-43 (A, B) and with 1 μM SQ-535 (C, D). Green channel settings (A, C): excitation 488 nm, signal collected from 515 to 600 nm. Red channel settings (B, D): excitation 560 nm, signal collected from 650 to 750 nm.

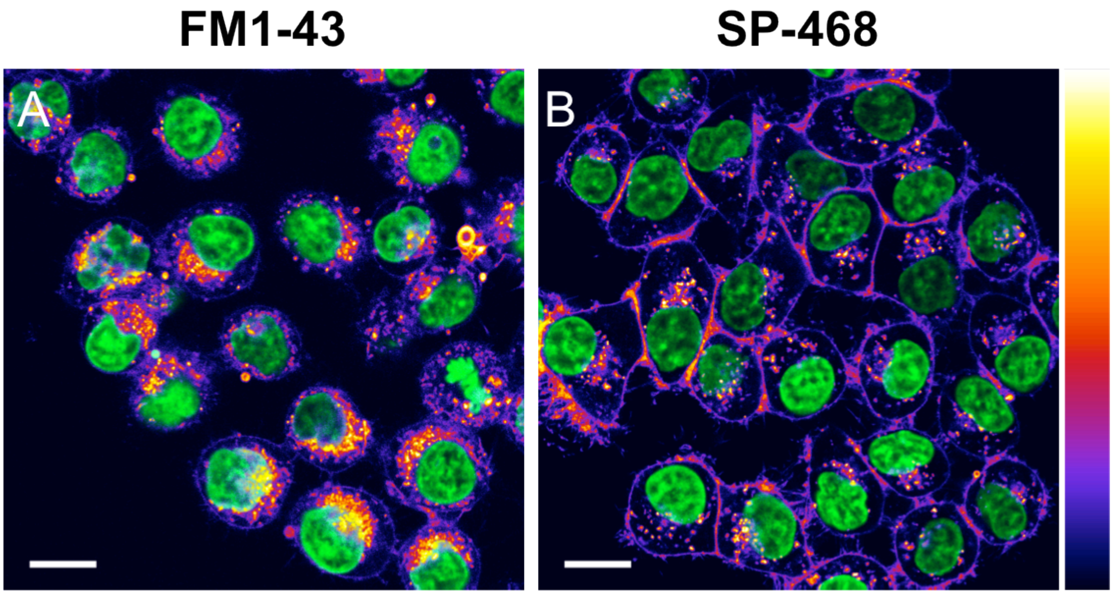


**Figure S8.** Laser scanning confocal imaging of live KB cells stained with 5 μM of FM1-43 (A) and 1 μM SP-468 (B) after 1h in Opti-MEM, depicting different pathways of internalization. Scale bar is 15 μm.
